# Insights into longevity and virus-driven adaptation from *Myotis* bat genomes

**DOI:** 10.1101/2024.10.10.617725

**Authors:** Juan M. Vazquez, M. Elise Lauterbur, Saba Mottaghinia, Léa Gaucherand, Sarah Maesen, Michael Singer, Sarah Villa, Melanie Bucci, Devaughn Fraser, Genavieve Gray-Sandoval, Zeinab R. Haidar, Melissa Han, William Kohler, Tanya M. Lama, Amandine Le Corf, Clara Loyer, Dakota McMillan, Stacy Li, Johnathan Lo, Carine Rey, Samantha L.R. Capel, Kathleen Slocum, Melissa Sui, William Thomas, Janet Debelak Tyburec, Rachel Brem, Richard Miller, Michael Buchalski, Jose Pablo Vazquez-Medina, Sébastien Pfeffer, Lucie Etienne, David Enard, Peter H. Sudmant

**Affiliations:** Department of Integrative Biology, University of California, Berkeley, Berkeley, CA USA; Department of Biology, Pennsylvania State University, State College, PA USA; Department of Ecology and Evolutionary Biology, University of Arizona, Tucson, AZ USA; Department of Biology, University of Vermont, Burlington, VT USA; Centre International de Recherche en Infectiologie (CIRI), Inserm U1111, Université Claude Bernard Lyon 1, CNRS UMR5308, Ecole Normale Supérieure ENS de Lyon, Université de Lyon, Lyon, France; Max Delbrück Center for Molecular Medicine in the Helmholtz Association, Robert-Rössle-Straße 10, Berlin, 13125, Germany; Université de Strasbourg, Architecture et Réactivité de l’ARN, Institut de Biologie Moléculaire et Cellulaire du CNRS, Strasbourg, France; Department of Molecular and Cellular Biology, University of California, Berkeley, Berkeley, CA USA; Wildlife Genetics Research Unit, Wildlife Health Laboratory, California Department of Fish and Wildlife, Sacramento, CA, USA; Wildlife Diversity Program, Wildlife Division, Connecticut Department of Energy and Environmental Protection, Burlington, CT, USA; Department of Biology, California State Polytechnic University, Humboldt, Arcata, CA USA; Department of Pathology and Clinical Laboratories, University of Michigan, Ann Arbor, MI USA; Department of Biological Sciences, Smith College, Northampton, MA USA; Department of Science and Biotechnology, Berkeley City College, Berkeley, CA USA; Center for Computational Biology, University of California, Berkeley, Berkeley, CA USA; Bat Conservation International, Austin, TX USA; Department of Ecology and Evolution, Stony Brook University, Stony Brook NY USA; Colossal Biosciences, Dallas, TX USA; Bat Survey Solutions, LLC, Tucson, AZ USA; Department of Plant and Microbial Biology, University of California, Berkeley, Berkeley, CA USA

**Keywords:** Aging, Bats, Cancer, Evolutionary Biology, Functional Genomics, Immunity, Infectious Disease

## Abstract

The genus *Myotis* is one of the largest clades of bats, and exhibits some of the most extreme variation in lifespans among mammals alongside unique adaptations to viral tolerance and immune defense^1–3^. To study the evolution of these phenotypes we generated cell lines and near-complete genome assemblies for eight closely related *Myotis* species. Using genome-wide screens of positive selection, analyses of structural variation, and functional experiments in primary cells, we identify patterns of adaptation contributing to longevity, cancer resistance, and viral interactions. We demonstrate distinct modes of adaptation to DNA and RNA viruses compared to all other mammals, with bats exhibiting genome-wide overrepresentation of positive selection for DNA virus-interacting proteins and elevated rates of copy number variation for RNA virus-interacting proteins. Characterization of *Myotis-*specific duplications of the key immune factor protein kinase R (*PKR*) reveals multiple ancient segregating trans-species copy number polymorphisms. We show that the recurrent evolution of longevity seen in *Myotis* is associated with positive selection in cancer pathways, and demonstrate a unique response to DNA damage in primary cells of the long-lived *M. lucifugus*. Together, our results suggest that bats’ remarkable longevity and immunity are linked through pleiotropic adaptations to viruses and aging-related disease.

## Introduction

Bats (order Chiroptera) represent approximately 20% of all known mammalian species and are one of the most phenotypically diverse clades of mammals^4^. Since their emergence 60 million years ago^5^, many bat lineages have independently evolved a wide variety of life history strategies and phenotypic traits, including exceptional longevity, viral tolerance, and immune defenses^2,3^. Systems in which shared traits have evolved *de novo* multiple times are powerful resources for dissecting the genetic basis of phenotypes. The largest genus of bats - *Myotis* - emerged approximately 33 million years ago^6^ and encompasses over 139 described species spanning six continents and a wide range of ecological niches^7^. *Myotis* species demonstrate some of the most extreme variation in lifespan amongst mammals^1,8^, including a six-fold difference in lifespan between the longest-lived species (*M. brandtii*, 42 years^9^, **Fig. 1A**) and the shortest-lived species (*M. nigricans*, 7 yrs^10^) which diverged approximately 10.6 million years ago^11^. In addition, *Myotis* species are representative of bats’ remarkable immune mechanisms which enable viral tolerance and pathogen resistance^12^ contributing to their role as key zoonotic reservoirs^2,13^.

**Fig. 1:**
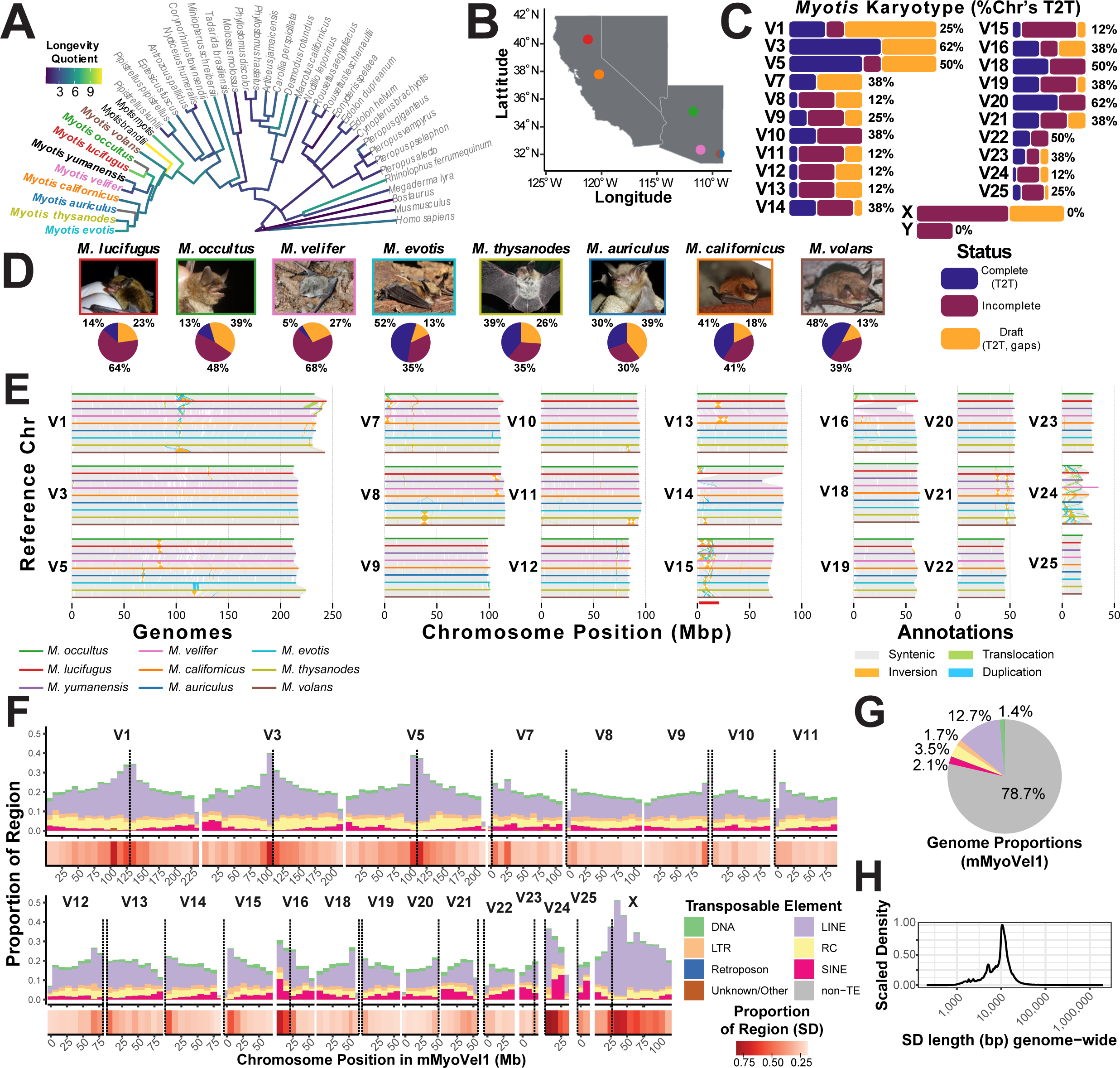
Near-complete reference assemblies reveals a varied structural variation landscape across 9 nearctic *Myotis* species. **A)** Phylogeny of Nearctic *Myotis* bats in this study, including outgroup species of bats, cow, mouse, and human. Branches are colored by their estimated *Longevity Quotient*, a ratio of observed-to-expected lifespan^1^. **B)** Map of capture sites in Arizona and California for samples generated in this study; each dot color indicates the species. **C)** Ideogram bar plot indicating completion status of each chromosome in assembly. Percentages next to ideograms indicate the proportion of T2T assembled chromosomes across species. **D)** Pie charts indicating the completion status of all chromosomes within each assembly, with representative images for the species shown above (see “Acknowledgements” for attributions). For **C-D**, “Complete (T2T)” status indicates that a chromosome is fully assembled telomere-to-telomere without gaps; “Draft (T2T, gaps)” status indicates that a chromosome is fully scaffolded with both telomeres, but has one or more gaps in the assembly; “Incomplete” status indicates that a chromosome was positively identified, but was not scaffolded from telomere to telomere (only contains one telomere). **E)** Synteny between chromosomes of 9 *Myotis* species showing syntenic regions (grey), inversions (orange), translocations (green), and duplications (blue). **F)** Distribution of transposable elements (top) and segmental duplications (red heatmap, bottom) in *M. velifer*. Putative locations of centromeres are denoted by the dotted lines. **G)** Pie chart of overall genomic proportions of TEs in *M. velifer*. **H)** Histogram of segmental duplication size distributions genome-wide in *M. velifer*.

To study how longevity and infectious disease resistance have evolved in *Myotis*, we used an integrated field-to-functional-genomics approach to assemble near-complete genome assemblies for eight *Myotis* species with primary cell culture resources for functional validation. We identify copy number variable genes associated with RNA viral tolerance and stress response as well as a trans-species copy number polymorphism of a key immune factor protein kinase R (*PKR*). Consistent with *Myotis*’ extreme lifespans relative to body size, we identified selective evolutionary signatures in genes associated with longevity- and cancer-related processes. In contrast to humans and other primates, in which virus adaptation is driven by interactions with RNA viruses, we find that modes of virus adaptation in bats differ between DNA and RNA viruses. Together, our results highlight pleiotropic adaptations contributing to the remarkable lifespan and immune phenotypes of *Myotis* bats.

## Results

### Eight near-complete *Myotis* genome assemblies

We collected skin punches and derived primary cell lines from several North American (“nearctic”)^14^ species (**Fig. 1A-B, D),** including from one of the longest-lived bats, *Myotis lucifugus*^15^. Using these cell lines and flash frozen tissues we generated *de novo* haplotype-resolved, chromosome-scale genome assemblies for eight species (**Fig. 1C-D**; **Extended Data Fig. 1**) using a combination of long-read PacBio HiFi sequencing and HiC scaffolding. These genomes are highly contiguous and near-complete, with an average of 98.6% (98.1-99%) of nucleotides assembled into the 22-23 syntenic^16^ chromosomal scaffolds; an average QV score of 66; and among the highest contig NG50s of any Chiropteran genome to date (**Extended Data Fig. 1**; **Supplementary Table 1**). We identified an average of 20,869 protein coding genes with mammalian homologs per genome, with BUSCO^17^ scores ranging from 98.2% to 98.5% (**Extended Data Fig. 1E**), which we used to build a time-calibrated, maximum likelihood tree of Chiroptera (**Extended Data Fig. 2A**; **Supplementary Table 2**). Across all eight genomes, each autosome has been completely assembled telomere-to-telomere (T2T) in at least one species (**Fig. 1C**); within assemblies, 29%-70% of chromosomes are fully assembled with an average of less than one gap per chromosome (**Fig. 1D; Supplementary Table 1**). Overall, these fully annotated genomes represent some of the most contiguous mammalian assemblies to date.

### Abundant structural variation in *Myotis*

We investigated the landscape of structural variation within the tightly-conserved *Myotis* karyotype. With only six exceptions across over 60 studied species, all *Myotis* have a conserved 2n=44 karyotype - a remarkable phenomenon for a genus spread across six continents and 33 million years of divergence^4,6^. The *Myotis* karyotype consists of three large autosomes; one small metacentric autosome; 17 small telocentric autosomes; and metacentric X and Y chromosomes^16^. Consistent with this broad cytological conservation, we find large scale synteny across the nearctic *Myotis* in this study (**Fig. 1E**). However, structural variants (SVs) including inversions, duplications, and translocations are relatively common within chromosomes, especially adjacent to putative centromeric regions (**Fig. 1E-F**). This includes a ∼20Mb block at the sub-telomeric end of Chr V15 that displays frequent and recurrent inversions and translocations across the nearctic *Myotis*, and which contains a 10Mb locus that was recently identified as a potential target of recent selection by adaptive introgression^18^ (**Extended Data Fig. 3D**).

Using SyRI^19^, we identified between 6,813 - 8,013 SVs per nearctic *Myotis* genome relative to their outgroup, *M. myotis*; 97 - 99% of these events were under 10Kb. In the three large autosomes, which constitute ∼30% of the genome, we cataloged an average of 509 SVs (**Supplementary Table 3**). In contrast, in the small autosomes, constituting ∼65% of the genome, we observed an average of 316 events, highlighting the distinct structural evolution between these chromosome types with SVs found at 2-fold higher density in larger autosomes (**Supplementary Table 3**). However, large (≥10Kb) duplications, inverted duplications, and inverted translocations were more common on small autosomes compared to the large autosomes (**Supplementary Table 3**). We also quantified the distribution of transposable elements (TEs) across chromosomes (**Fig. 1F, G**). Consistent with prior observations in other Chiropteran genomes^20^, LINE elements appeared to be enriched around the predicted centromere (**Fig. 1F**) with simulations showing that this trend is primarily limited to the long, metacentric autosomes (**Extended Data Fig. 3A**; **Supplementary Table 3**). The concentration of segmental duplications was also significantly correlated with TE density in each species (linear regression, p < 0.001; **Fig. 1F, H**; **Extended Data Fig. 3B**) highlighting the importance of TEs in facilitating structural evolution. Together these results highlight high levels of structural variation occurring in the context of a highly constrained karyotype.

### A trans-species *PKR* copy number variant

Among the structural variants identified in nearctic *Myotis*, the gene Protein Kinase R (*PKR*/*EIF2AK2*) - previously shown to be duplicated in certain *Myotis*^21^ - stood out as an interferon-stimulated gene with antiviral activity against both DNA and RNA viruses. Using our near-complete genome assemblies, we fully resolved the sequence and structure of the two previously known haplotypes: H1, containing a single copy of *PKR* (*PKR2*); and H2, containing two tandemly duplicated copies of *PKR* (*PKR1* and *PKR2*; **Fig. 2A**). We also identified a third haplotype - H3 - with three tandem duplicates of *PKR* (*PKR1*, *PKR2*, and a third copy) present only in *M. californicus*. Remarkably, while 7 out of 9 *Myotis* species carried duplicated haplotypes, 5 of these cases were heterozygous for the duplicated haplotype (i.e. H1/H2 or H2/H3; **Fig. 2B**). Two *Myotis* individuals (*M. lucifugus* and *M. evotis*) only carried non-duplicated haplotypes (i.e. H1/H1; **Fig. 2B**). To determine the evolutionary history of the duplicates, we used GeneRax^22^ to construct a tree from alignments of all *PKR* gene copies across Neartic *Myotis*, using *Pipistrellus pygmaeus* as a non-*Myotis* outgroup (**Fig. 2C**). We find that *PKR2* is the ancestral copy of *PKR*, and that *PKR1* originated from a single duplication event at the root of *Myotis*. These results highlight that both the duplicated and unduplicated haplotypes have likely been segregating for tens of millions of years, representing an ancient trans-species polymorphism.

**Fig. 2:**
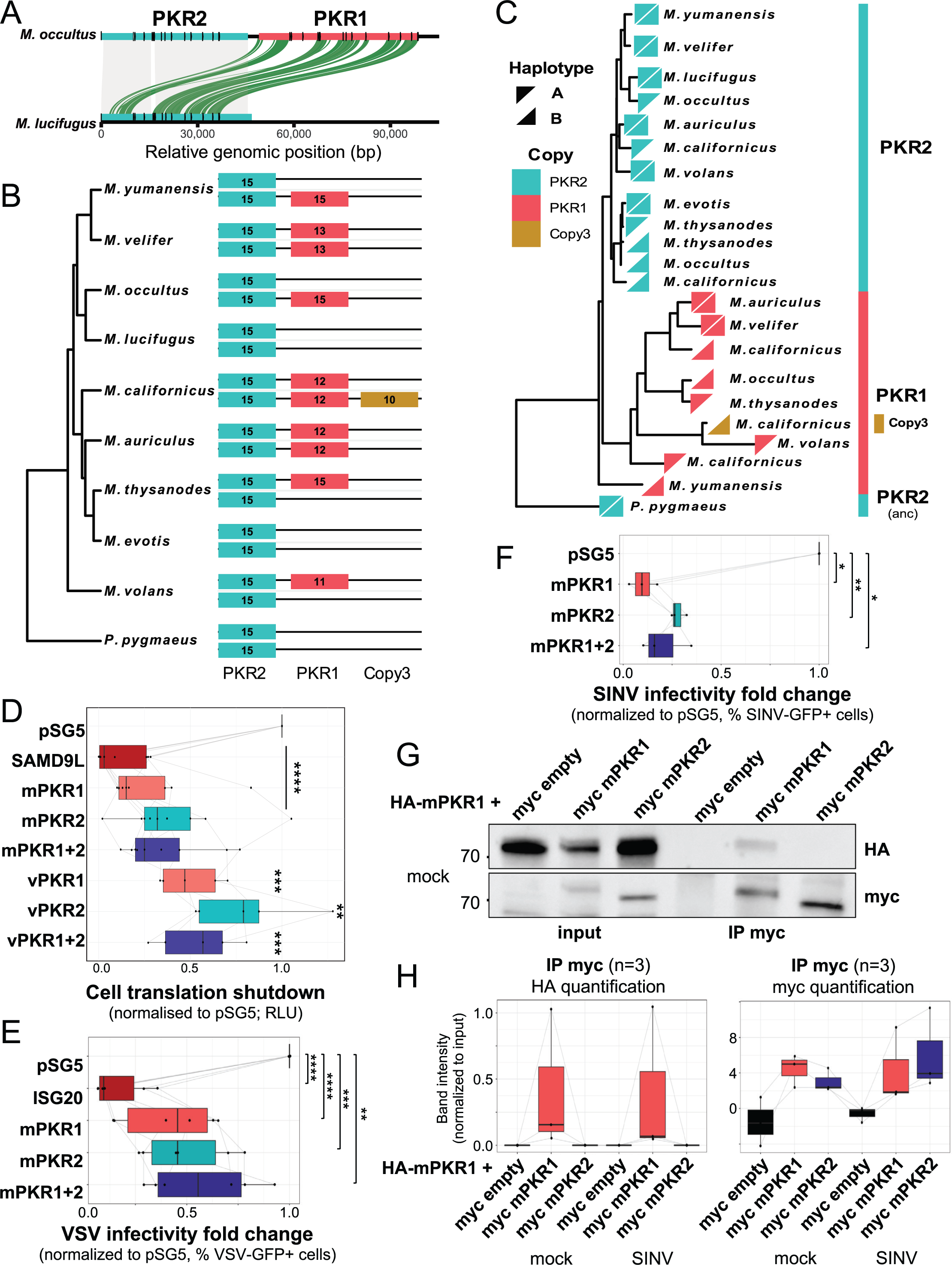
Evolutionary history and function of an actively segregating copy number polymorphism of *PKR* in *Myotis*. **A)** Structural comparison of PKR haplotypes in two species. Orthologous regions are indicated by grey bands, syntenic duplications are indicated in green, and exons with black marks. **B)** Cartoon of the PKR locus in the two phased haplotype assemblies of each *Myotis* species in this study, with the number of exons per copy. While *PKR2* is present across all haplotypes, *PKR1* and *PKR copy 3* are polymorphic. **C)** Reconciled gene tree for *PKRs* across all haplotypes and species shown in B. Haplotype corresponding to the reference (A) and alternate (B) haplotype for each species are represented by upper- and lower-diagonal triangles, respectively. **D)** Effect of *Myotis* PKR expression on luciferase reporter translation in PKR-KO cells, measured in Relative Light Units (RLU) and normalized to the empty pSG5 control. Human SAMD9L-GoF is a positive control of translation inhibition^72^. **E-F)** Effect of PKR on viral infection by VSV-GFP (E, and S3B) and SINV-GFP (F 34h post-infection, and S3C), measured by flow cytometry as the % of GFP+ cells, normalized to the pSG5 control. Although all conditions restricted VSV and SINV, expression of both PKR1+PKR2 was not beneficial against these viral infections. ISG20 served as a positive control of VSV-GFP restriction^71^. **G)** Co-immunoprecipitation (IP) of PKR-KO cells transfected with *Myotis* HA-PKR1 and *Myotis* myc-PKR1, *Myotis* myc-PKR2, or myc-empty vector control. Proteins were pulled down with anti-myc beads and lysates from 5% input or IP samples were run on a western blot and stained for HA and myc (see **Extended Data** Fig. 4C for SINV conditions). **H)** Quantification of the three independent experiments (mock and SINV) for HA (left) and myc (right). Error bars indicate mean ± SEM for at least three independent experiments. Statistical significance was assessed using an unpaired t-test (*, p<0.05; **, p<0.01; ***, p<0.001; ****, p<0.0001). mPKR indicates *Myotis myotis* PKR; vPKR indicates *Myotis velifer* PKR. For gel source data, see **Supplementary** Figure 1.

PKR is a stress response and innate immune factor that becomes active upon detection of dsRNA through sensing, auto-phosphorylation and dimerization^23^. Active PKR shuts down protein translation and restricts viral replication^23^. Although PKR1 and PKR2 are both expressed in basal or interferon-stimulated *Myotis velifer* cells, only their independent functional impacts have previously been investigated^21^. Given the co-expression and the complex haplotype diversity we identified in *Myotis*, we set out to determine whether the two *PKR* copies act additively, synergistically, or as dominant negatives in several essential functions. We investigated the functional impact of the duplicates’ co-expression on total PKR protein expression levels, cell viability, inhibition of translation, antiviral restriction and homo/hetero-dimer formation (**Fig. 2D-H**). To pinpoint the contribution of PKRs, we used a heterologous cell system^21^ with PKR-KO HeLa cells transfected with either an empty pSG5 plasmid (control), or pSG5 encoding PKR1, PKR2, or PKR1+PKR2 (1:1 ratio) from *Myotis myotis* (myoMyo) or *Myotis velifer* (myoVel). In this system, PKR transfection leads to its autoactivation^21^.

First, we found that, although *Myotis* Flag-PKR1 was expressed at lower levels than Flag-PKR2, PKR1+PKR2 co-expression did not affect the overall PKRs’ steady-state protein expression levels (**Extended Data Fig. 4A**). Second, we performed cell viability assays at two doses of transfected PKRs and found that cell viability was only affected at high dose (**Extended Data Fig. 4D**). Third, using a luciferase reporter translational assay, we found that PKR1+PKR2 coexpression led to an intermediate translation shutdown, showing neither synergistic nor dominant negative effect of the PKR duplicates (**Fig. 2D**). Fourth, we tested PKR1+PKR2 co-expression effects on infections by two different model RNA viruses: VSV-GFP (Vesicular stomatitis virus encoding a GFP reporter, a representative of *Rhabdoviridae* family that is common in bats, including *Myotis*) and SINV-GFP (Sindbis virus, a representative of *Togaviridae*) (**Fig. 2E-F**; **Extended Data Fig. 4C, E**). We found that PKR1+PKR2 restricted viral infections to a similar extent – or slightly lower in the case of VSV – as PKR1 or PKR2 alone (**Fig. 2E-F**; **Extended Data Fig. 4C, E**, p>0.05). While PKR1 homodimerized, PKR1 and PKR2 did not form heterodimers (**Fig. 2G-H**; **Extended Data Fig. 4B** in SINV condition). This suggests that, since their duplication, divergence between *Myotis* PKR copies has been particularly strong at the protein-protein interface hindering their capacity to form heterodimers.

These results suggest that PKR1 and PKR2 do not exhibit dominant-negative or synergistic effects but instead act additively in their primary cellular effector functions. While their duplication may broaden resistance to certain DNA and RNA viral antagonists^21^, the increased cell toxicity observed at high PKR doses suggests a tradeoff that could explain the absence of PKR duplication in other mammals. This tradeoff may also account for the persistence of both duplicated and non-duplicated haplotypes in *Myotis*. Consistent with this, PKR1, which is generally more potent in our experiments, is expressed at significantly lower levels than PKR2 in *Myotis velifer* cells^21^. Together, these results reveal gene duplicates still segregating across several *Myotis* species, likely shaped by a balance between antiviral defense and cell toxicity.

### Modes of adaptation to DNA & RNA viruses

To further explore how viruses may have shaped the genomes of *Myotis* bats we tested for adaptive signatures in virus-interacting proteins (VIPs) in *Myotis* and other bats. VIPs are host proteins that physically interact with viral proteins (e.g. *CD45*, **Fig. 3A**), and can be proviral (contributing to viral infection, e.g. viral receptors), antiviral (protective against viral infection, e.g. interferons), or both depending on infection stage and virus type. VIPs commonly have major roles in basic host function, and as a result most host cell biological pathways include VIPs. Previous studies investigating positive selection across mammals have found an enrichment for adaptation among a set of 5,527 manually curated VIPs, defined as host proteins that have at least one experimentally verified physical interaction with a viral protein, RNA, or DNA^24^.

**Fig. 3:**
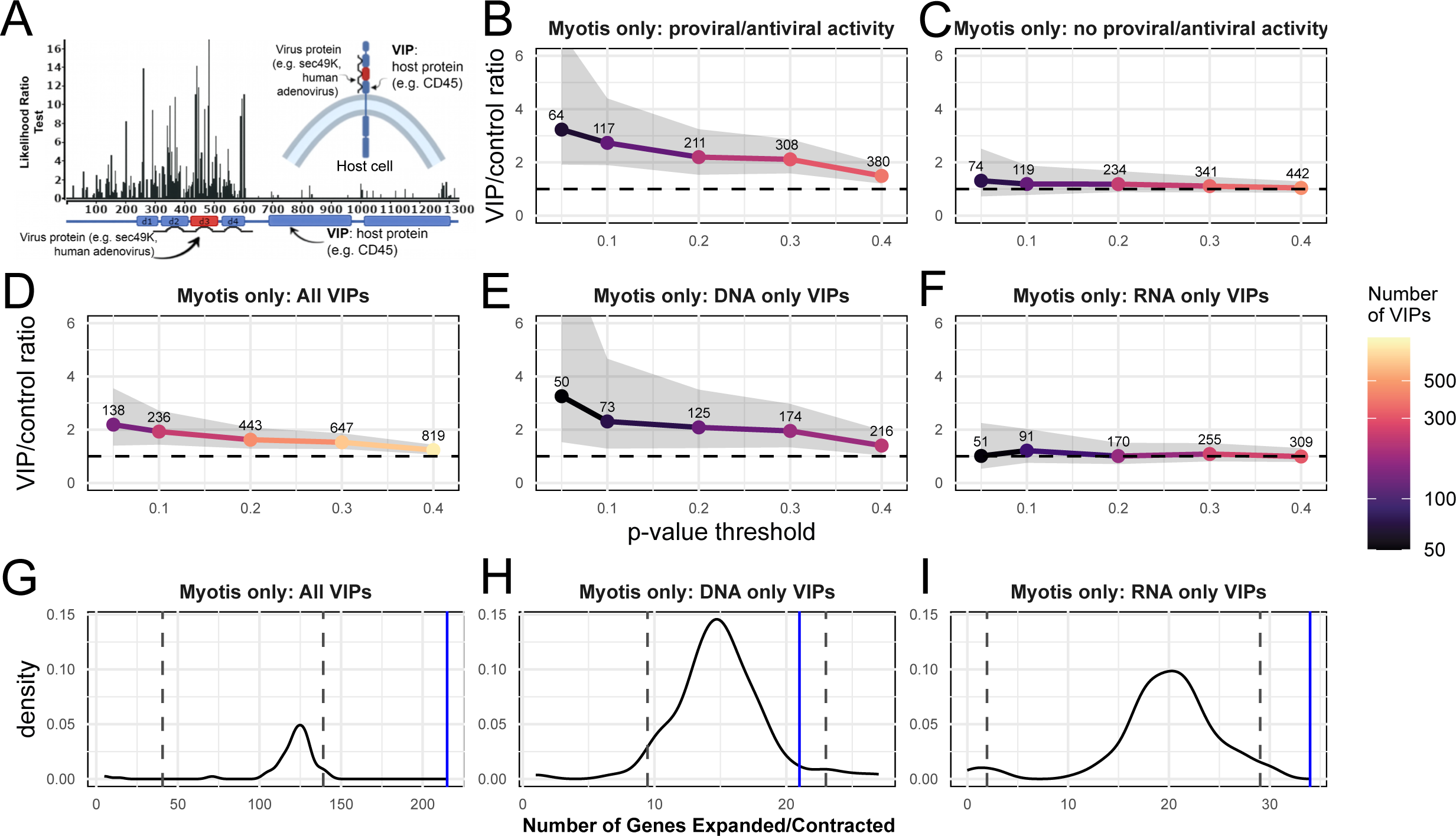
Adaptation to DNA viruses, but not RNA viruses, is enriched in *Myotis* and other bats. **A)** Diagram of an example VIP, host cell transmembrane receptor CD45, showing strength of adaptation at each codon and the locations of direct interactions with the human adenovirus protein sec49K. **B-F)** Enrichment plots showing the ratio of positive selection in only *Myotis* bats in VIPs versus matched sets of control genes at different p-value thresholds. The solid line shows the median ratio; the color of the line and the number above each point represent the number of VIPs with significant BUSTED-MH p-values at the given threshold, and therefore used with the same number of matched control genes for each ratio calculation; the grey band represents the 95% confidence interval generated by bootstrapping sets of matched control genes. The 95% confidence interval widens at lower p-values because there are fewer VIPs with significant p-values as the alpha threshold decreases, reducing power. **G-I)** The number of VIP genes inferred by CAFE to have significantly expanded or contracted in copy number on at least one branch of the nearctic *Myotis* phylogeny (blue line) compared to the distribution across 100 randomized sets of non-VIP control genes (black line). Vertical dashed lines indicated the 95% highest density interval for the control gene distribution.

By calculating an enrichment score from the ratio of positive selection in VIPs compared to their matched control genes using BUSTED-MH^25^, we found that, like other mammals, *Myotis* show an enrichment for adaptation at VIPs (**Fig. 3B-D**; **Supplementary Table 5**). Physical host-virus interactions may not always result in fitness effects in the host. We therefore repeated our analysis using a gene set restricted to VIPs with experimental evidence of specific pro- or anti-viral effects, and thus with a stronger expectation of fitness effects. We expect that if viruses were not the primary driver of adaptation in VIPs, then there would be no difference in the enrichment of selection in VIPs with versus without demonstrated pro- or anti-viral effects. However, we observed an even stronger significant elevation in the ratio of positive selection in these proviral and antiviral VIPs (**Fig. 3B**; **Supplementary Table 5**), but no elevation in this ratio in VIPs without experimental evidence of specific pro- or anti-viral effects (**Fig. 3C**; **Supplementary Table 5**). This is consistent with the expectation of viral interaction as the cause of enrichment of positive selection in VIPs in bats^26^. We repeated this analysis using a dataset of 47 publicly-available non-*Myotis* bat genomes, and confirmed these same patterns across bats more broadly, even when excluding *Myotis* genomes (**Extended Data Fig. 5A**).

Previous work has suggested that bats may have different physiological responses to DNA and RNA viruses^27^. To determine if this was reflected in genomic VIP adaptation, we compared the enrichment of positive selection in VIPs that interact only with DNA viruses (DNA-only VIPs) to those that interact only with RNA viruses (RNA-only VIPs). Remarkably, we found that VIP adaptation in *Myotis* and other bats is driven by selection only in DNA VIPs (**Fig. 3E**; **Extended Data Fig. 5B**) in marked contrast to the observed pattern in RNA VIPs, which show no evidence of genome-wide enrichment in adaptation (**Fig. 3F**; **Extended Data Fig. 5C**).

Adaptation to viral pathogens can also occur via gene copy number changes^28–30^. We tested whether VIPs were enriched among genes that were recently gained or lost throughout *Myotis* using the method of Huang *et al*. (2023) to test the cumulative per gene, per branch copy number changes and birth-death rates of VIPs compared to other non-VIP genes.^31^ While the birth-death rates of VIP and non-VIP genes were comparable (p = 0.071; **Extended Data Fig. 5D**), VIP genes were significantly more likely to have undergone expansions and/or contractions on at least one branch of the *Myotis* family (p < 0.001; **Fig. 3G**). Furthermore, we find that this pattern is driven exclusively by copy number changes in RNA-only VIPs, and not by changes in copy number of DNA-only VIPs (**Fig. 3H-I**). This suggests that, relative to the general variation in gene family birth-death rates across species, RNA VIPs are more dynamic across the nearctic *Myotis* as a whole.

In contrast to what we observe in bats, VIP adaptation in humans is driven by positive selection in RNA - and not DNA - VIPs^32^. To investigate if DNA VIP-driven adaptation in bats is exceptional among mammals, we replicated these analyses across four other large mammalian clades that are well represented among publicly-available mammalian genomes: Primates, Glires, Euungulata, and Carnivora. We found that while other mammalian orders show a mix of adaptation enrichments in both RNA and DNA VIPs, none show the absence of genome-wide enrichment of protein-coding adaptation in RNA VIPs observed in bats (**Extended Data Fig. 5A-C**). These results highlight that bats, including *Myotis*, exhibit unique modes of adaptation to DNA viruses compared to RNA viruses, in contrast to all other mammals.

### Body size & lifespan evolution in bats

Beyond their remarkable immune adaptations, bats are exceptional as the longest-lived clade of mammals after correcting for their body size^1^, demonstrating an ∼11x range of lifespans within a ∼650x-range of body sizes^8,33^. *Myotis* species in particular exhibit the full dynamic range of bat lifespans within a narrow range of body sizes. While bats have been noted as an exception to the strong allometric scaling (positive correlation with body size) of lifespan otherwise seen across mammals and other metazoans, this exception has not been tested using phylogenetically corrected statistics leveraging well-resolved phylogenies.

To test the hypothesis of non-allometric scaling of lifespan in bats, we modeled the evolution of body size and lifespan independently across a supertree of over 1000 mammals^34^ (**Fig. 4A-B**; **Extended Data Fig. 6A-B**; **Supplementary Table 6**). We generally observed agreement between evolutionary patterns of body size and lifespan such as in whales (Cetacea)^35^, elephantids (Proboscidea)^36^, and primates^37^ (**Fig. 4A-B**). However, while only minor changes were observed in their body size, we observed some of the largest and most-rapid changes in lifespan across mammals in bats (**Fig. 4A-B**; **Extended Data Fig. 6A-B**; **Supplementary Table 6**). These changes are largely independent between species and genera, consistent with the theory of multiple independent increases in lifespan across bats. We used phylogenetically-corrected generalized linear models and ANCOVA to quantify the relationship between body size and lifespan across mammals. We find that bats experience a 40% greater increase in lifespan per 1% increase in body size compared to non-bat mammals (0.223% increase in lifespan versus 0.159% increase in lifespan per 1% increase in mass); these rates, however, were not significantly different after phylogenetic correction (**Extended Data Fig. 6C-D**; pANCOVA: p=0.29). In *Myotis*, we saw many of the fastest increases in lifespan relative to their most recent ancestor, as measured by the change in their lifespan over the divergence time from their most recent ancestor (“Δlifespan”): *Myotis brandtii* (8.6 Δlifespan, 99th percentile), *Myotis lucifugus* (8.4 Δlifespan, 99th percentile), *Myotis myotis* (2.5 Δlifespan, 93nd percentile), *Myotis grisescens* (1.3 Δlifespan, 92nd percentile), and the *Myotis* common ancestor (1.7 Δlifespan, 89th percentile) (**Fig. 4B**; **Extended Data Fig. 6B; Supplementary Table 6**). Together these results demonstrate that *Myotis* bats and their recent ancestors have evolved some of the most extreme increases in lifespan among mammals, despite similar allometric lifespan scaling of bats and mammals after phylogenetic correction.

**Fig. 4:**
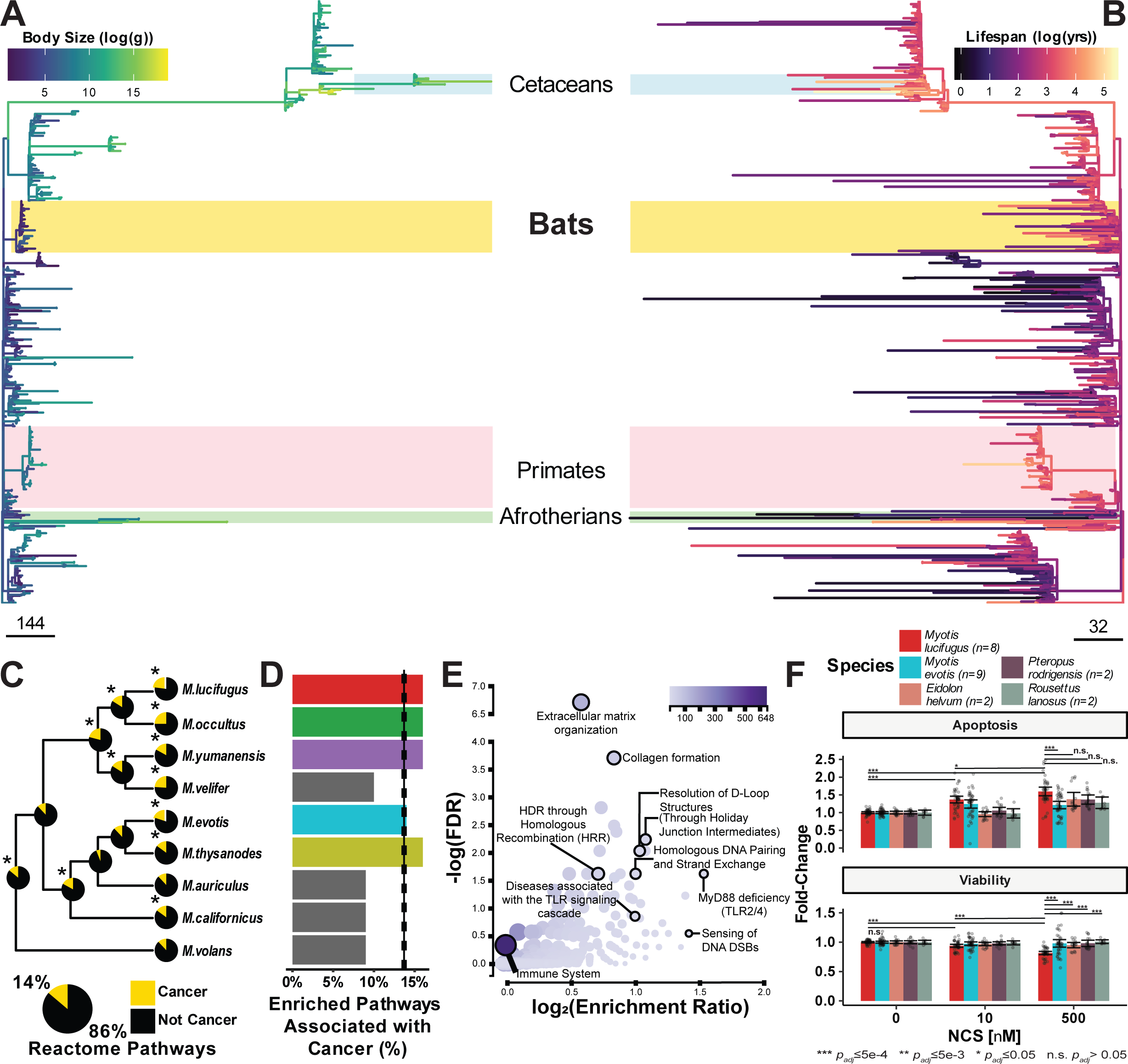
Evolution of cancer risk and resistance in *Myotis* and other mammals. **A, B)** Cophylo plot of the evolution of body size (**A**) and lifespan (**B**) across *Eutheria*. Branch lengths in **A-B** are scaled proportional to the rate of change of the trait over time. **C)** The phylogeny of nearctic *Myotis* species with inset pie charts representing the proportion of the top 100 Reactome pathways overrepresented among genes under selection at each node that are associated with cancer-related processes. Below, pie chart indicates expected proportion of pathways are cancer-associated after 1000 random samples of 100 pathways from the full Reactome database. Asterisks represent nodes with proportions greater than the expected value at p≤0.05 using Fisher’s exact test. **D)** Proportion of the top 100 Reactome pathways overrepresented among genes under selection across all nodes in a species’ evolutionary history that are associated with cancer-related processes. **E)** Volcano plot of overrepresented pathways in Reactome among the union set of genes under selection across all nodes in the evolutionary history for *M. lucifugus*. **F)** Viability and Apoptosis fold-change in 5 bat species in response to different doses of neocarzinostatin (NCS), a potent inducer of DNA double-strand breaks. Points represent individual replicates normalized to each species’ control, while bars represent mean ± 95% confidence intervals.

### Pleiotropic immune & DNA damage response

Rapid changes in body size and lifespan can have major implications for the evolution of cancer risk and resistance across mammals^38^. While lifetime cancer risk scales proportionally to both body size and lifespan *within species*^38,39^, there is little to no correlation between body size, lifespan, and cancer risk *across species*^39,40^. This observation, which has been coined *Peto’s Paradox*^40^, suggests that species with more cells or longer lifespans have adapted to reduce their cancer risks.

We hypothesized that the extreme changes in lifespan observed throughout the *Myotis* phylogeny should exert a selective pressure on genes associated with cancer processes. Using aBSREL^41^ to test for selection at terminal and internal branches within *Myotis*, we found that, per node, an average of 5.46% and 20.8% of protein-coding genes had at least one region under positive or negative selection, respectively (**Supplementary Table 7**). Genes under either positive or negative selection were enriched for several pathways in immunity, cancer, and aging, with many intersecting multiple of these processes, suggesting possible pleiotropic selective pressures (**Supplementary Table 7**).

We quantified the proportion of cancer-associated pathways^42^ overrepresented among genes under positive selection throughout the phylogeny (**Fig. 4C-D**). Among genes under positive selection, we observed that most nodes within nearctic *Myotis* were enriched for cancer hallmark pathways, especially at the recent ancestors of the longest-lived species (e.g. *M. lucifugus* + *M. occultus*; **Fig. 4C**). Furthermore, when considering the union of all genes under positive selection in the evolutionary history of each species since their common ancestor, we observed significant enrichments in the representation of cancer-associated pathways in many of the lineages with the greatest cumulative increases in lifespan (*M. lucifugus*, *M. occultus, M. evotis, M. yumanensis;* **Fig. 4D; Extended Data Fig. 6E**).

The longest-lived bat in our study, *M. lucifugus*, had an overrepresentation of pathways specifically associated with DNA double-strand break (DSB) repair when looking at both lineage-wide and node-specific enrichments in positive selection using the Reactome database^43^ (**Fig. 4E**; **Supplementary Table 7**). This includes 35 out of 65 genes in the high-fidelity Homologous Recombination Repair pathway, and 21/37 members of the Homology-Directed Repair via Single Strand Annealing (**Fig. 4E**; **Extended Data Fig. 6E**; **Supplementary Table 7**). These results suggested that *M. lucifugus* might have an enhanced response to DNA DSBs relative to other bats. To test this hypothesis, we assessed the tolerance of *M. lucifugus* to neocarzinostatin (NCS), a potent radiomimetic agent that induces DNA double-strand breaks^44^ (**Fig. 4F**), compared to *M. evotis* and three non-*Myotis* bats (*Eidolon helvum, Pteropus rodrigensis,* and *Rousettus lanosus*). At low doses of neocarzinostatin, *M. lucifugus* was the only species with sensitivity to neocarzinostatin after 24 hours, with a drop in viability and concomitant increase in apoptosis. At high doses, *M. lucifugus* had the highest level of apoptosis and the greatest drop in viability of all the bats tested. This is consistent with other long-lived species, including elephants^45,46^, naked mole rats^47^, and bowhead whales^48^, where longevity is associated with an increased ability to clear out damaged cells.

To better understand the genes that may be driving this difference, we examined RNA expression in *M. lucifugus* cells after 6 and 18 hours of treatment with 100 nM NCS. At 6 hours we observed up-regulation of genes associated with cell cycle arrest (e.g. *MDM2*, *CDKN1A*) and cell death (e.g. *BAX*), and down-regulation of genes in pathways associated with cell division, growth, and DNA damage repair and synthesis. At 18 hours, we observe up-regulation of genes in pathways associated with cellular stress response and continued down-regulation of the cell cycle (**Fig. 5A-D**; **Extended Data Fig. 7A**; **Supplementary Table 9**). However, we observed that the strongest pathway enrichment in our DNA-damage RNA-seq data were genes associated with the innate immune response, including genes in the PI3K/AKT pathway (**Fig. 5C-D**; **Supplementary Table 9**). We hypothesized that the observed relationship between the unique DNA damage response of *M. lucifugus* may be due to pleiotropy with DNA VIPs. Indeed, we observed that there was a highly significant enrichment for genes that are differentially expressed after NCS treatment and DNA-only VIPs (p=3.34e-5, **Fig. 5E**; α=0.0245, **Extended Data Fig. 7B**). The genes at the intersection of DNA-only VIPs and NCS response are overrepresented in pathways associated with the cell cycle, senescence, transcription, and DNA damage repair (**Fig. 5F**). We repeated these analyses only considering genes under positive selection in *Myotis lucifugus*. The intersection between DNA-only VIPs and NCS-DE genes remained highly significant (p=1.88e-3, **Extended Data Fig. 7C**), with only the pathways “Cell Cycle”, “DNA Damage Repair”, and “Deubiquination” enriched at FDR≤0.05 (**Fig. 5G**). DNA-only VIP genes which are not NCS-responsive did not show these signatures (**Extended Data Fig. 7D-G).** Together these results suggest that selection on the innate immune response may drive *agonistic* pleiotropy in aging-associated traits like DNA damage response and cell cycle regulation.

**Fig. 5:**
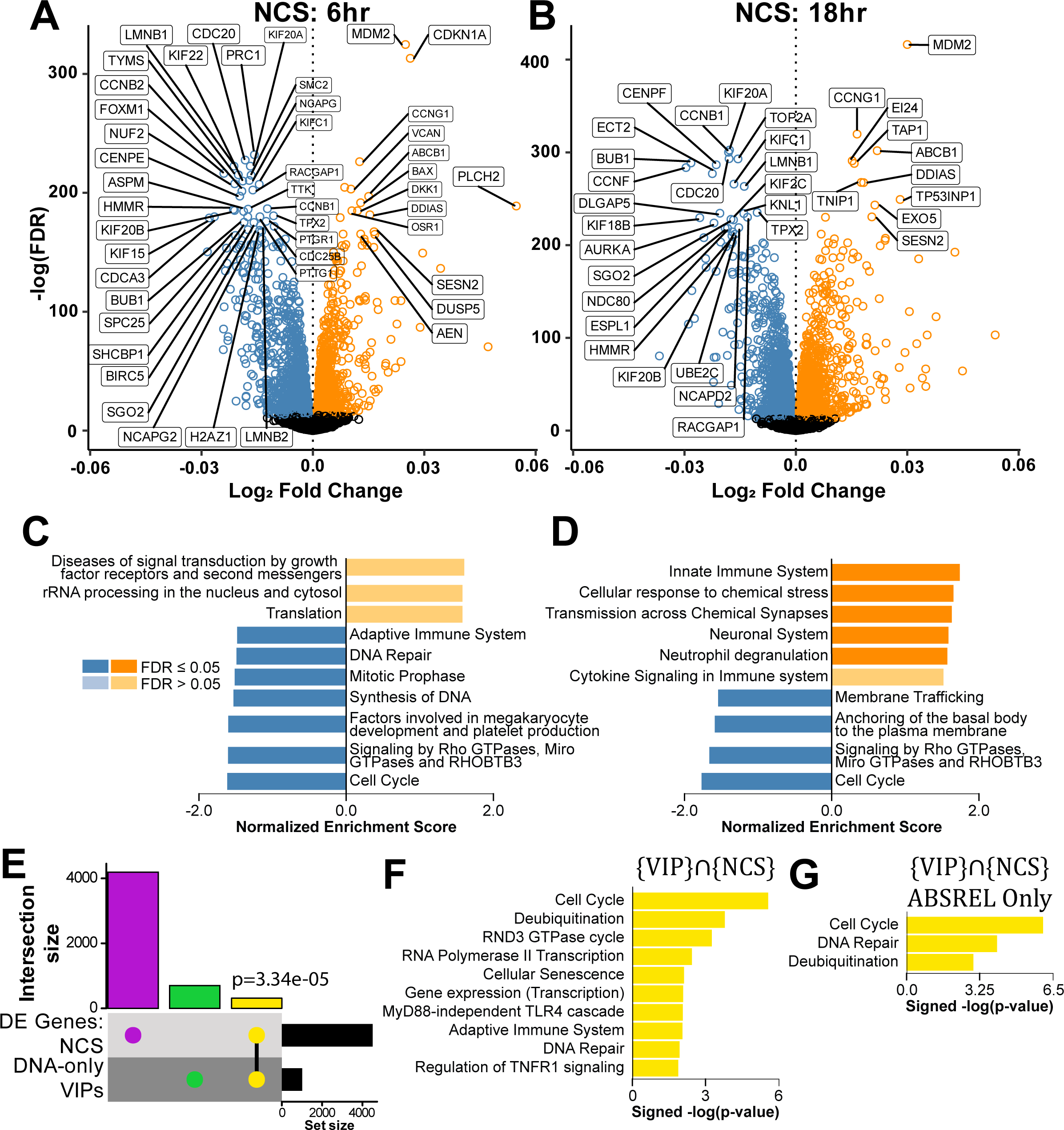
Pleiotropy among genes under selection in *M. lucifugus* between DNA-damage responsive genes and DNA-only VIPs. **A-B)** Volcano plots of genes that are differentially expressed in *M. lucifugus* after 6 hours (**A**) and 18 hours (**B**) of treatment with 100nM neocarzinostatin (NCS). **C-D)** Pathway-level Gene Set Enrichment Analysis (GSEA) of genes differentially expressed at 6 and 18 hours post-treatment. **E)** Upset plot between genes that are differentially expressed after NCS treatment and DNA-only VIPs; the hypergeometric P-value of the intersection is shown. **F)** Pathway-level overrepresentation analysis of genes that are DNA-only VIPs and are also differentially expressed in response to NCS treatment. **G)** Pathway-level overrepresentation analysis of genes identified as under selection in *M. lucifugus* via aBSREL that are DNA-only VIPs and are also differentially expressed in response to NCS treatment.

## Discussion

In addition to evolving true flight, bats are remarkable for their long lifespan^8,33^, stress tolerance^39,49^, and viral tolerance^2,50^. The genes and mechanisms underlying these highly complex and pleiotropic phenotypes can be challenging to identify, especially for rapidly-evolving phenotypes such as host-pathogen interactions. We identify new patterns of adaptation contributing to longevity, cancer resistance, and viral interactions in bats, and demonstrate a unique DNA damage response in primary cells of the long-lived *M. lucifugus*.

The evolution of body size and lifespan across mammals has major implications for the co-evolution of cancer risk and resistance. We find altered scaling of longevity in *Myotis* that is predicted to have serious consequences for their intrinsic, per-cell cancer risk. Similar to other systems where the evolution of cancer resistance has been largely driven by rapid changes in body size^36,51,52^, the rapid and repeated changes in lifespan across an order of magnitude observed in *Myotis* are hypothesized to result in strong selective pressures on lifetime cancer suppression to avoid malignancies^33,38,53^. This is supported both by recent meta-analyses on cancer risk in vertebrates^39,54^ demonstrating little to no correlation between neoplasias and either species body size or lifespan; as well as by studies showing weak or no correlation between mutation rates and either species body size or lifespan, respectively^55–57^.

Bats are also important models for understanding infectious disease response and immune adaptation because of their tolerance to viruses and role as zoonotic reservoirs^13,27,58^. Here, we show that while bats have adapted to both DNA and RNA viruses, they have done so by different modes - protein evolution versus copy number changes, respectively. This is in contrast to humans and other primates, in which protein coding adaptations to RNA viruses dominate^32,59^. In addition, we demonstrate complex patterns of structural variation in immune genes, including a segregating duplication of *PKR*, which encodes a major protein involved in the antiviral innate immune system. *PKR* has functional relevance in its activity against both DNA and RNA viruses^21^. We further demonstrate the functional implications of copy number variation in *PKR*, including an additive effect on cellular effector functions.

Multiple hypotheses have been proposed to connect the particular physiology and ecology of bats with the evolution of remarkable adaptations such as viral infection tolerance and defense, stress tolerance, and exceptional longevity^3^. Many theories of the driving causes of disease resistance and longevity in bats have been proposed, including the evolution of flight^1,60–62^, the disposable soma hypothesis^63^; metabolic state^64^; torpor^8^; and other environmentally-driven adaptations^65,66^. Additionally, many studies have highlighted the intersection of these traits, including links between hibernation, DNA repair, longevity^8,58^, and infectious disease resistance^8,27,58,67^. Our results support the importance of *agonistic* pleiotropy in shaping bats’ evolution, wherein genetic adaptations for many specific traits (e.g. DNA virus innate immunity) may prove beneficial to other seemingly-unrelated traits (e.g. cancer resistance, cellular homeostasis, longevity). Importantly, bats are also the only known mammal to harbor active DNA transposons^68^, the suppression of which may also contribute to signatures of selection in DNA-interacting VIPs. Our findings on pervasive selection on DNA-only VIPs and the extreme evolution of longevity-associated in *Myotis* and other bats support a broader hypothesis that traits such as cancer risk, cellular homeostasis, and antiviral response have evolved in tandem due to pleiotropic selection at coinciding points in bats’ evolutionary history.

## Supporting information

Supplementary Information Guide

## Acknowledgements

Analyses were performed using the following High Performance Computing (HPC) resources hosted by the following organizations: the University of Arizona (supported by the University of Arizona TRIF, UITS, and Research, Innovation, and Impact (RII); maintained by the UArizona Research Technologies department); the University of California, Berkeley (Savio HPC, supported by the UC Berkeley Chancellor, Vice Chancellor for Research, and Chief Information Officer); and the University of Vermont (Vermont Advanced Computing Center). PacBio HiFi sequencing was done by the DNA Technologies and Expression Analysis Cores at the UC Davis Genome Center (NIH SIG 1S10OD010786-01). We acknowledge the contribution of the SFR Biosciences (UAR3444/CNRS, US8/Inserm, ENS de Lyon, UCBL) ANIRA cytometry platform, especially Véronique Barateau and Estelle Devevre. We thank Adam Geballe (Fred Hutch Cancer Center) for sharing Hela PKR-KO cells. We thank the members of the LP2L team (CIRI) and members of the Bat1K consortium for helpful discussions. As co-contributors, M.E.L. and J.M.V. reserve the right to order their names interchangeably when citing this work.

Image attributions for the bat photos featured in Fig. 1: U.S. National Park Service (*M. auriculus, M. thysanodes*); ; Juan Cruzado Cortés (*M. californicus, M. volans, M. occultus*); Ansil B.R. (*M. velifer*); Issac Krone (*M. evotis*); and SMBishop & the Wikimedia Foundation (*M. lucifugus*).

## Author Contributions

Conceptualization, J.M.V., M.E.L., D.E., P.H.S., L.E., D.F., M.B.; Methodology, J.M.V., M.E.L., D.E., P.H.S., L.E., G.G.S.; Software, J.M.V., M.E.L., D.E., P.H.S., L.E., T.M.L.; Validation, J.M.V., M.E.L., D.E., P.H.S., L.E., J.P.V.M.; Formal Analysis, J.M.V., M.E.L., D.E., P.H.S., L.E., S.M., A.L.C., D.M., S.V.; Investigation, J.M.V., M.E.L., S.M., M.B., D.F., G.G.S., L.G., Z.R.H., M.H., W.K., T.M.L., A.L.C., C.L. S.M., D.M., S.L., J.L., C.R., S.L.R.C., M.S., K.S., W.T., J.D.T., S.V., R.M., M.B., J.P.V.M., S.P., L.E., D.E., P.H.S.; Resources, J.M.V., M.E.L., D.E., P.H.S., L.E., M.B., D.F., G.G.S., L.G., Z.R.H., M.H., W.K., T.M.L., A.L.C., S.M., D.M., S.L., J.L., C.R., S.L.R.C., M.S., K.S., W.T., J.D.T., S.V., R.M., M.B.; Data Curation, J.M.V., M.E.L., S.M., M.B., D.F., G.G.S., L.G., Z.R.H., M.H., W.K., T.M.L., A.L.C., S.M., D.M., S.L., J.L., C.R., S.L.R.C., M.S., K.S., W.T., J.D.T., S.V., R.M., M.B., J.P.V.M., S.P., L.E., D.E., P.H.S.; Writing - Original Draft, J.M.V., M.E.L., D.E., P.H.S., L.E., S.M., J.L.; Writing - Review & Editing, J.M.V., M.E.L., S.M., M.B., D.F., G.G.S., L.G., Z.R.H., M.H., W.K., T.M.L., A.L.C., S.M., D.M., S.L., J.L., C.R., S.L.R.C., M.S., K.S., W.T., J.D.T., S.V., R.M., M.B., J.P.V.M., S.P., L.E., D.E., P.H.S.; Visualization, J.M.V., M.E.L., D.E., P.H.S., L.E., S.M., A.L.C.; Supervision, J.M.V., M.E.L., D.E., P.H.S., L.E.; Project Administration, J.M.V., M.E.L., D.E., P.H.S., L.E.; Funding Acquisition, J.M.V., M.E.L., D.E., P.H.S., L.E.

## Additional Information

Supplementary Information is available for this paper. Correspondence and requests for materials should be addressed to P.H.S. or L.E. for all PKR-related work. Reprints and permissions information is available at www.nature.com/reprints.

## Materials and Methods

### Data availability

All sequencing data and genomes generated in this study are available on NCBI under Bioprojects PRJNA973719, PRJNA1035541, and PRJNA1295526. Annotations generated in this study are available at https://github.com/docmanny/myotis-gene-annotations. All other data are archived on DataDryad at DOI: 10.5061/dryad.02v6wwqfh.

### Code availability

All custom code used herein is available at https://github.com/sudmantlab/MyotisGenomeAssembly.

### Samples and materials

All sample collection efforts were performed following approved Animal Use Protocols by the University California, Berkeley and the University of Arizona. All samples collected in this study were collected under Scientific Collection Permits from either the Arizona Game and Fish Department (SP405504, SP407155, SP403977, and SP407113) or the California Department of Fish and Wildlife (S-220230001-22025-001) (see **Supplementary Table 1**). Bats were sampled using standard mist-netting procedures, including taking standard body measurements, following USGS recommendations for White-Nose Syndrome and COVID-19 prevention^69,70^. For *M. lucifugus*, the donor individual was field-caught in California and transported to the Genetics Laboratory of the California Department of Fish and Wildlife, where they were euthanized via isofluorane. The *M. velifer* individual was caught in Arizona and euthanized in the field via isoflurane. For both *M. lucifugus* and *M. velifer*, tissues were collected and preserved via flash-freezing in liquid nitrogen. All other genomes were generated from primary cell lines. Additional information can be found in the *SI Guide* and in **Supplementary Table 1**.

All PKR experiments were performed using HeLa PKR-KO cells (kindly provided by A. Geballe, Fred Hutchinson Cancer Center, Seattle WA). The cells were maintained at 37°C with 5% CO and cultured in DMEM supplemented with 5% fetal bovine serum (FBS), 1% penicillin/ streptomycin mix and 1 μg/mL puromycin (Sigma-Aldrich). All transfections were performed 24 hours after seeding, using 3 µL of TransIT-LT1 Transfection Reagent (Mirus Bio) per 1 µg of DNA and Opti-MEM media. We used previously-generated pSG5-FLAGx2 vectors encoding either *M. myotis* PKR1 (GenBank OP006550), *M. myotis* PKR2 (GenBank OP006559), *M. velifer* PKR1 (GenBank OP006558), or *M. velifer* PKR2 (GenBank OP006557)^21^. Plasmids encoding the interferon-stimulated gene ISG20^71^ and a constitutively active variant of the sterile alpha motif domain-containing protein 9-like SAMD9L-F886Lfs*11 (here, SAMD9L^72^) were used as controls in viral infections and cell translation experiments, respectively.

### Near-complete genome assembly and annotation

Details on genome assembly, including DNA and RNA extraction, preparation of PacBio HiFi, Omni-C (Dovetail Genomics), and RNA-seq libraries, genome assembly, annotation, and manual curation can be found in the *SI Guide*.

### Structural Variation

To understand the genomic distribution of structural variants, including segmental duplication events, we used SyRI (Synteny and Rearrangement Identifier^19^). After masking repetitive regions such as telomeres and centromeres, the primary 22 scaffolds corresponding to the autosomes of the nearctic *Myotis* genomes were mapped to each other in the correct orientations using minimap2^73^. We ran SyRI on the resulting files and plotted the results with plotsr^74^.

### Phylogenetics

A phylogeny of all 536 mammals in our alignments was generated using IQTREE^75^ (version 2.3.1) using all gene alignments with the settings “-B 1000 -m GTR+F3×4+R6.” Gene trees were generated from gene alignments to exclude alignments with less than 50% gaps in the sequence and 4 or more species represented by using IQTREE with settings “--wbtl --bnni --alrt 1000 -B 1000 --safe”. The best substitution models for each gene were saved as a NEXUS file. As the bootstrap values for this tree were unanimously 100%, we ran ASTRAL-IV^76^ (v1.24.4.7) using the settings “-t 54 -u 2 -C” and confirmed that our phylogeny agreed with other previously-published *Eutherian* phylogenies^77,78,79^. The Chiroptera portion of our phylogeny was time-calibrated using MCMCtree^80^ and PAML^81^ (v. 4.10.0) with the bat-subset of our codon alignments and using fossil calibrations (see Supplementary Table **2**)^78,82–90^. We ran MCMCtree twice to generate the Hessian matrix and confirm convergence, and ran 10 independent chains using the “out.BV” file from the first run. Finally, the output files of all 10 chains were combined to compute final divergence time estimates.

### Ancestral Body Size and Lifespan reconstruction

To explore how body size and lifespan have evolved over time in mammals, we used a super-phylogeny of mammal species^34^ subsampled to only contain species with extant body size and lifespan data collected from AnAge^15^ and PanTHERIA^91^. Ancestral body sizes and lifespans were simulated separately using StableTraits^92^ and the settings “--iterations 100000 --skip 1000 --chains 8”. Estimates were further validated by comparing our initial results to a second run of StableTraits using the same settings.

### Selection Scans & Evolutionary Rates

#### aBSREL

To conservatively test for branch-specific selection, we used aBSREL^41,93^ (version 2.5.48) to test for selection at each branch within the nearctic *Myotis* clade for 15,734 gene alignments spanning 536 mammals. These genes were identified as 1:1 orthologs across the full alignment, with no more than 50% sequence gaps and at least 4 species present in the alignment. For each gene alignment, the full phylogeny was trimmed to match the species present, and the HyPhy script “label-tree.bf” was used to highlight all nodes within nearctic *Myotis* as foreground branches. aBSREL was then run using the pruned and highlighted tree and gene alignments on the command line with the options “--code Universal --branches Foreground". We defined a gene as under selection if one or more regions within the gene demonstrated a selective pressure with an FDR-corrected p-value ≤ 0.05. A gene was specifically identified as under positive selection if at least one of these significant regions had an ⍵>1.

#### BUSTED

To quantify the total amount of positive selection across the *Myotis* tree or the different species trees used in this manuscript, we used an improved version of the BUSTED^25,93^ test called BUSTED-MH (see *SI Guide* for details on BUSTED vs BUSTED-MH). We applied BUSTED-MH to 19,646 *Myotis* orthologous CDS alignments with at least five orthologs. These orthologs are cases where the Orthofinder gene trees coincide with the species tree. This effectively removes issues regarding whether we should use the gene or the species tree, at the cost of removing 2,110 genes from the *Myotis* selection analysis. Similarly, we applied BUSTED-MH to 17,469 non-*Myotis* bat alignments with at least five orthologs. The species in these two sets (*Myotis* bats and non-*Myotis* bats) are mutually exclusive. This includes a subset of 14,091 alignments with orthologs present in two thirds of the non-*Myotis* bat species, regardless of the genes’ presence/absence in the *Myotis*-only complement. This is specifically used to show that patterns of virus-driven adaptation are representative of all bats. We also tested 17,890 primate alignments with at least five orthologs with BUSTED-MH, as well as 19,311 glire, 18,000 carnivora and 18,504 ungulate alignments. Similarly, HyPhy MEME (with – resample 100 to increase power) was run to identify individual codon sites subject to diversifying selection in the *CD45* gene for visualization with host protein-virus contact sites.

#### Enrichment scores for VIPs

We expanded on the set of VIPs reported in Souilmi *et al*. (2021)^26^ (5,291 VIPs) by searching the literature for more recent publications reporting VIPs, with a final dataset of 5,527 VIPs. These VIPs (“All VIPs”) were further sub-categorized into 4 groups based on their interactions with either DNA viruses and/or RNA viruses: “DNA VIPs”, “DNA-only VIPs”, “RNA VIPs”, and “RNA-only VIPs” (**Supplementary Table 5**). For each VIP set, we selected sets of matching control genes by controlling for 16 different. Sets of control genes were resampled in a bootstrap procedure to generate 95% confidence intervals for sets of genes across a range of p-values; additional details can be found in the *SI Guide*.

#### Gene Duplications and CAFE analysis

To quantify patterns of gene duplication and loss, we quantified the copy number of genes with human orthologs from our gene annotations for each nearctic *Myotis* genome. To calculate per-gene expansion and loss rates and their statistical significance, we ran CAFE^94^ v5 on the previously described set of copy number counts using our time-calibrated species tree pruned to include only the nine nearctic *Myotis* species. CAFE uses a birth-death model of gene family evolution to investigate changes in gene family size accounting for evolutionary history using the species phylogeny. Evolutionary branch length is incorporated into the model expectations, thus accounting for potential bias resulting from differences in branch lengths. We ran CAFE on the subset of genes with two or more copies in at least one species using a Poisson distribution for the root frequency (*-p*), first generating an error model to correct for genome assembly and annotation error (*-e*). We compared the base model (each gene family belongs to the same evolutionary rate category) to gamma models (each gene family can belong to one of *k* evolutionary rate categories) with different values of *k* from 2 to 13. A final gamma model with *k*=9 was chosen to balance model log likelihood with the number of gene families for which the optimizer failed. The model was run three separate times to ensure convergence.

To understand if genes in these pathways have higher birth-death rates or are more likely to have significant changes in gene copy number than expected relative to other genes, we evaluated the gene copy birth-death rate LJ and number of genes that have significantly expanded or contracted in copy number on at least one branch within our nearctic *Myotis* phylogeny using a bootstrap procedure. Following Huang et al.^31^, we tested if VIP genes in particular underwent significant copy number changes or had significantly different birth-death rates than non-VIP genes. For each category of VIP genes (all VIPs, DNA VIPs, DNA only VIPs, RNA VIPs, and RNA only VIPs), we generated 100 bootstrap sets of control non-VIP genes with the same number of genes as the corresponding VIP gene set. We ran CAFE on each set of VIP genes and the corresponding control non-VIP genes to infer per-gene birth-death rates and per-gene, per-branch expansion/loss events.

### Assessment of DNA Double-Strand Break Tolerance

We assessed each species’ tolerance to DNA double strand breaks using a by measuring viability, cytotoxicity, and apoptosis across a range of doses of Neocarzinostatin, a radiomimetic drug. We measured dose response curves in wing-derived primary dermal fibroblasts across 5 bat species (*Myotis lucifugus*, n=8; *Myotis evotis*, n=8; *Rousettus langosus*, n=2; *Eidolon helvum*, n=2; *Pteropus rodrigensis*, n=2) using the multiplexed ApoTox-Glo assay (Promega). Using 96-well plates, two individuals and 11 doses were assessed simultaneously with four technical replicates. Results were normalized to treatment controls for each individual in R (see code in repository).

### PKR1/2 Characterization in PKR-KO HeLa cells

#### Western blot

Details on PKR Western blots can be found in the *SI Guide*.

#### Luciferase reporter assays

Details on PKR luciferase reporter assays can be found in the *SI Guide*.

#### PKR co-immunoprecipitation

HeLa PKR-KO cells were transfected with 1.25 µg plenti6 HA-tagged *M. myotis* PKR1 plasmid per million cells and 1.25 µg of either plenti6 myc empty vector, myc-tagged *M. myotis* PKR1 or myc-tagged *M. myotis* PKR2 plasmid using TransIT-LTI transfection reagent (Mirus Bio). The next day, some wells were infected with Sindbis virus expressing GFP (SINV-GFP) at MOI 2 for 24 hours. Cells were then scraped with cold PBS and pelleted. For the IP, cells were lysed in 500 µl IP buffer (50 mM Tris HCl pH 7.5, 140 mM NaCl, 6 mM MgCl2, 0.1% NP40) supplemented with RNase (RiboLock, Fisher Scientific) and protease (complete EDTA-free protease inhibitor cocktail, Sigma) inhibitors for 10 minutes on ice, then centrifuged at 12,000 *xg* for 10 minutes at 4° C. 5% of the volume was kept for input, while the rest was incubated with 40 µl µMACS anti-c-myc MicroBeads (Miltenyi Biotec) for 1 hour at 4° C with constant rotation. Samples were then loaded onto µMACS columns placed in the magnetic field of a µMACS Separator (Miltenyi Biotec), washed 4 times with cold IP buffer, and eluted with the µMACS denaturing elution buffer. Proteins were denatured in elution buffer for 5 minutes at 95° C, then loaded onto a 4-20% BioRad Criterion TGX Stain-Free precast gel and transferred onto an Amersham Protran nitrocellulose membrane (Sigma) for 1 hour. Membranes were blocked for 1 hour in 5% milk in PBS (Euromedex) supplemented with 0.2% tween (Fisher) and incubated with mouse anti-myc monoclonal antibodies (Abcam 9E10, cat# ab32) then secondary anti-mouse IgG antibodies conjugated with HRP (Sigma, cat# A4416), or with rat anti-HA antibodies conjugated with HRP (Roche, Sigma, cat# 12013819001). Images were taken on a Fusion FX imager (Vilber) with SuperSignal West Femto Chemiluminescent Substrate (ThermoFisher Scientific).

#### Cell viability assay

HeLa PKR-KO cells were transfected 24 hours after plating in 96 well plates (n=10,000 cells/well), with 100 or 200 ng of pSG5 plasmid: empty or coding for *M. myotis* or *M. velifer* PKR1, PKR2 or PKR1+2 equal mix (50%-50%). 24 hours post-transfection, positive control cells were treated with an apoptosis-inducing drug, Etoposide, at different doses (250, 200 or 100 µM). 48 hours post transfection, cells were harvested and lysed to quantify luminescent signal according to CellTiter-Glo Luminescent Cell Viability Assay (Promega) kit protocol.

#### VSV and SINV infections

VSV infections: 200,000 cells were plated in 12-well and transfected 24 hours after with 350 ng of pSG5 plasmid: empty, or encoding *M. myotis* or *M. velifer* PKR1, PKR2, or equal input of PKR1 and PKR2 (175 ng per plasmid), or a plasmid encoding interferon-stimulated exonuclease gene 20 (ISG20), due to its known antiviral activity against VSV as positive control^71^. Cells were infected 24 hours post transfection with replicative VSV-GFP virus^95^ at a MOI of 3. Cells were fixed with 4% paraformaldehyde 16-18 hours post infection. VSV infection was quantified by measuring the percentage of GFP positive cell populations with BD FACSCanto II Flow Cytometer (SFR BioSciences). Fold change results were normalized to the empty pSG5 condition across at least three independent experiments.

SINV infections: HeLa PKR-KO cells were transfected with 5 µg pSG5 empty vector, *M. myotis* PKR1, *M. myotis* PKR2 or 2.5 µg *M. myotis* PKR1 + 2.5 µg *M. myotis* PKR2 per million cells using TransIT-LTI transfection reagent (Mirus Bio). The next day, some wells were infected with SINV-GFP at MOI 0.2. Cells were then placed into a CellCyte X live cell imaging system (Cytena) and pictures of every well were taken every 2 hours for 48 hours. The fraction of GFP+ cells over the total cell area was measured and averaged from six photos of 2 individual wells per condition, and repeated for a total of three independent experiments.

## Extended Data Legends

**Extended Data Fig. 1: Contiguity statistics for 8 novel *Myotis* genomes. A)** NG-decay plot for contig-level assemblies of bat genomes used in this paper. **B)** LG-decay plot for contig-level assemblies of bat genomes used in this paper. **C)** NG-decay plot for scaffold-level assemblies of bat genomes used in this paper. **D)** LG-decay plot for scaffold-level assemblies of bat genomes used in this paper. The invariance of the curves in **A-D** corresponding to the genomes generated in this paper is due to the high-contiguity of contigs assembled by *hifiasm-hic*; chromosome-level scaffolds were largely unaffected by scaffolding efforts. **E)** Mammalian BUSCO scores for annotations generated for the 8 new *Myotis* genomes. **F-M)** Genome assembly metric comparisons between assemblies derived from tissues or from primary cell lines. While assembly metrics do not correlate with coverage of PacBio HiFi sequencing (**F**), assemblies derived from primary cell lines do show improved statistics.

**Extended Data Fig. 2: Estimated node divergence times for Chiroptera and ASTRAL-IV scores. A)** Distribution of time estimates for nodes across the full fossil-calibrated Chiroptera phylogeny. **B)** ASTRAL-IV scored phylogeny of all mammalian genomes used in this study. **C)** ASTRAL-IV scored phylogeny of Chiroptera. For **B-C**, the local posterior probability (LPP) scores of nodes under <90% are shown on the tree; the branch at nearctic *Myotis* root with low bootstrap support is highlighted in red in **A**.

**Extended Data Fig. 3: Correlations of TEs and structural variation in *Myotis* genomes. A)** Histogram of LINE element counts in 1Mb bins across chromosomes in *M. velifer*. Black lines indicate the estimated location of the centromeric region as a 20Mb bin. **B)** Correlation between the proportion of bases within segmental duplications and the proportion of bases within TEs in bins across 9 Nearctic *Myotis* genomes. The blue line represents the linear regression of the correlation between the two proportions, and the R^2^, p-value, and equation for the correlations are plotted in each graph. **C)** Correlation between the proportion of bases within segmental duplications and the proportion of bases within TEs in bins for *M. velifer* per TE family**. D)** Synteny between chromosome V15 across 9 *Myotis* species. The binned distribution of transposable elements (top) and segmental duplications (red heatmap, bottom) in *M. velifer* are shown above the syntenic alignment in phylogenetic order. The ∼20-Mb block at the subtelomeric end of Chr V15 spans several immune-related and interleukin signaling genes, including IL-1 and IL-36.

**Extended Data Fig. 4: Additional functional characterization of *Myotis* PKR duplicates. A)** Steady-state protein expression levels in PKR-KO HeLa cells expressing FLAG-tagged *Myotis myotis* PKR1 and/or PKR2. Western blots targeting FLAG or Tubulin (loading control) in lysate of cells transfected with either empty vector (pSG5), PKR1-FLAG, PKR2-FLAG, or an equimass mix of both PKR vectors (total 350 ng or 700 ng). **B)** Co-immunoprecipitation (coIP) of PKR-KO HeLa cells transfected with plasmids encoding HA-tagged *Myotis myotis* PKR1 and either myc-tagged *Myotis myotis* PKR1, myc-tagged *Myotis myotis* PKR2 or a myc empty vector control in SINV-GFP infected conditions. Proteins were pulled down with anti-myc coated beads and lysates from 5% input or immunoprecipitated samples were run on a western blot and stained for HA and myc. Image representative of 3 independent experiments. **C)** PKR-KO HeLa cells transfected with plasmids encoding an empty vector control, *Myotis myotis* PKR1, PKR2 or both PKR1 and PKR2 were infected with SINV-GFP at MOI 0.2 and GFP fluorescence was monitored over time by live imaging. Mean +/- standard deviation of the percentage of GFP fluorescent area over total cell area was plotted for 3 independent experiments, each representing the average value of six photos from two duplicate wells. **D)** Effect of PKRs on cell viability, normalized to the control. Etoposide treatments are positive controls of cell death. Only high transfection of PKRs impacted cell viability (200 ng total of DNA for 10,000 cells). mPKR, *Myotis myotis* PKRs; vPKR, *Myotis velifer* PKRs. Error bars indicate mean ± SEM for at least three independent experiments. Statistical significance was assessed using an unpaired t-test (*, p<0.05; **, p<0.01; ***, p<0.001; ****, p<0.0001). **E)** Effect of *Myotis velifer* PKRs on viral infection by VSV-GFP, measured by flow cytometry as the % of GFP+ cells, normalized to the pSG5 control. ISG20 served as a positive control of VSV-GFP restriction^71^. For gel source data, see **Supplementary Figure 1**.

**Extended Data Fig. 5: VIP selection enrichments in bats and other mammals. A)** Enrichment for selection of all VIPs (left), DNA-only VIPs (middle), and RNA-only VIPs (right) versus matched control genes in non-*Myotis* bats. **B)** Enrichment for selection of DNA-only VIPs versus matched control genes in 4 non-bat clades of mammals. **C)** Enrichment for selection of RNA-only VIPs versus matched control genes in 4 non-bat clades of mammals. For **A-C**, plots show the ratio of selection (□) p-values of viral interacting proteins (VIPs) v.s. matched controls for VIPs and control gene sets identified as under positive selection by BUSTED-MH as a function of p-value threshold. **D)** CAFE p-value estimates for the birth-death rate (λ) of VIP genes and VIP-subsets compared to randomized matched sets of non-VIP genes. The 95% confidence interval of the distribution of the bootstrapped non-target control genes is indicated by the dotted lines around each distribution; the p-value estimated for the target gene sets are indicated by the solid blue line.

**Extended Data Fig. 6: Evolution of body size and lifespan across *Myotis*. A-B)** Estimates of body size (**A**), and lifespan (**B**) in Chiroptera with major bat families highlighted. **C-D)** Phylogenetic least-squares regression for lifespan (log(yrs)) as a function of body size (log(kg)) across mammals (**C**) and bats (**D**) in the Upham *et al*. (2019)^34^ phylogeny; lines represent the phylogenetically-corrected generalized least squares regression for each group indicated in the legend. **E)** Raincloud plots of genes under selection (p_adj_≤0.05) by aBSREL in each terminal species. Each histogram represents the distribution of the highest omega (□) value for each gene after multiple testing correction (p_adj_≤0.05). The 95% confidence interval and median for significant □’s are represented by the black bar and circle, respectively; the 95% confidence interval and median □ for all genes are shown in grey below. Individual genes’ □’s are represented by colored points. Genes that are associated with pathways previously explored in the literature for bats, such as insulin signaling, iron metabolism, DNA damage response, and SERPIN-family genes, are highlighted.

**Extended Data Fig. 7: Genes differentially expressed after neocarzinostatin treatment and their overlap with DNA-only VIPs**. **A)** Heatmap of log_2_ Counts Per Million (CPM) for the top 50 differentially expressed genes in *M. lucifugus* after 6- and 18-hours of treatment with 100 nM neocarzinostatin. **B)** Histogram of p-values for the overlap of 10,000 simulated sets of non-NCS DE genes and non-DNA-only VIPs. The empirical false positive rate, □, is 0.0245. **C)** Upset plot of genes that are either 1) under selection (*ABSREL*, p_adj_≤0.05) in *Myotis lucifugus* ({*aBSREL*}); 2) differentially-expressed after neocarzinostatin treatment ({*NCS*}); or 3) are classified as DNA-only VIPs ({*VIP*}). The hypergeometric p-value for the set {*NCS*}∩{*aBSREL*}∩{*VIP*} is shown. **D-G**) Pathway-level overrepresentation analysis of genes that are exclusive to either {*NCS*} (**D, E**) or {*VIP*} (**F, G**) either before (**D, F**) or after (**E, G**) sub-settings for only those genes identified as under selection. Only pathways significant at FDR≤0.05 are shown.

**Supplementary Data Table 1.** Genome statistics, fossil data, and MCMCtree settings for time tree

**Supplementary Data Table 2.** Accessions for additional genomes used in this study

**Supplementary Data Table 3.** SyRI-identified structural variants (SVs)

**Supplementary Data Table 4.** Experimental data for PKR experiments **Supplementary Data Table 5.** List of VIPs and VIP subclasses

**Supplementary Data Table 6.** Ancestral state reconstructions, pANCOVA statistics for body size and lifespan in mammals

**Supplementary Data Table 7.** aBSREL significant gene lists and Reactome enrichments

**Supplementary Data Table 8.** Experimental data for Neocarzinostatin experiments

**Supplementary Data Table 9.** RNA-seq QC statistics and GSEA results

