## Supplementary figures and images for "Insights into longevity and virus-driven adaptation from *Myotis* bat genomes"

### ExtendedDataFig_1.pdf

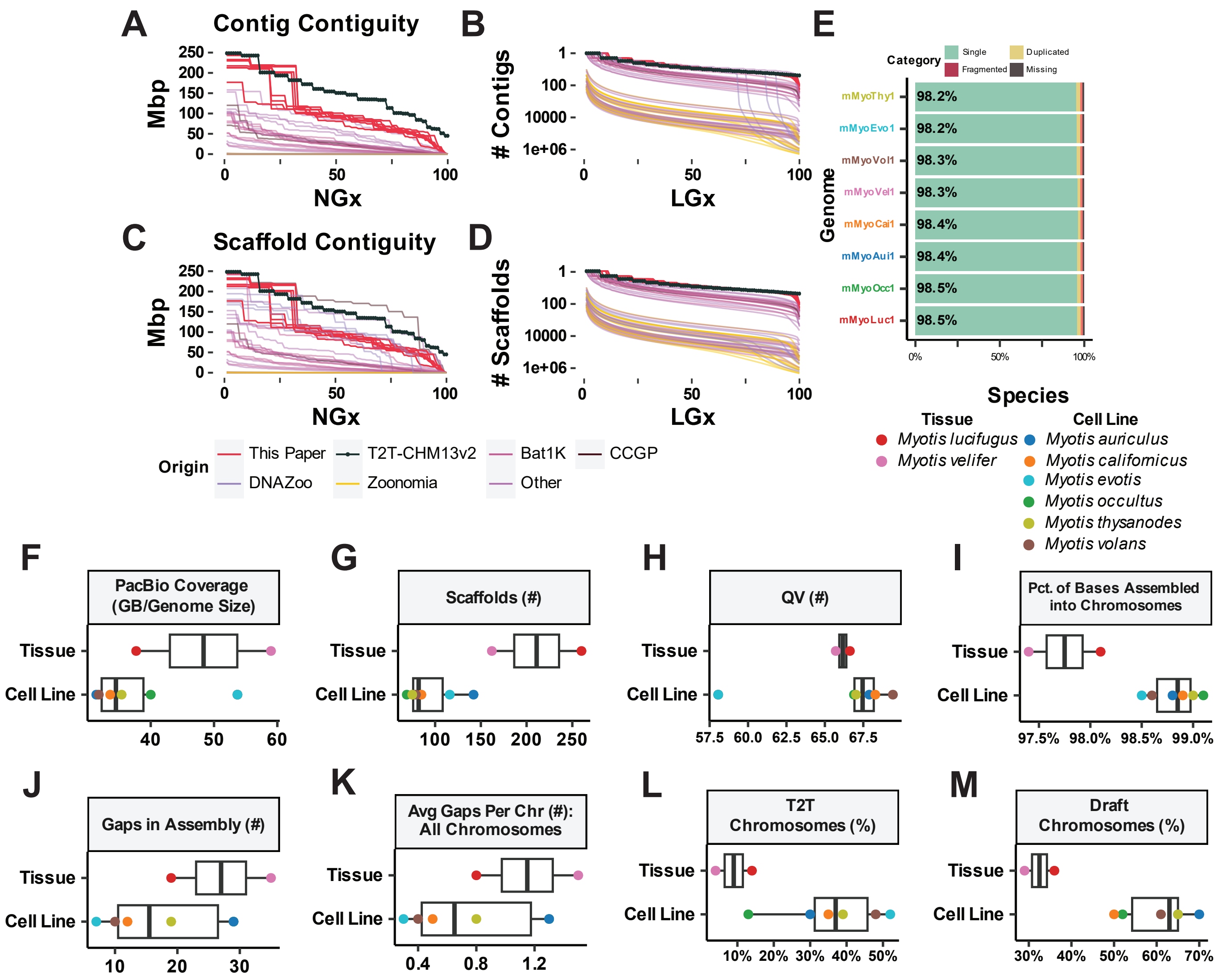

### ExtendedDataFig_2.pdf

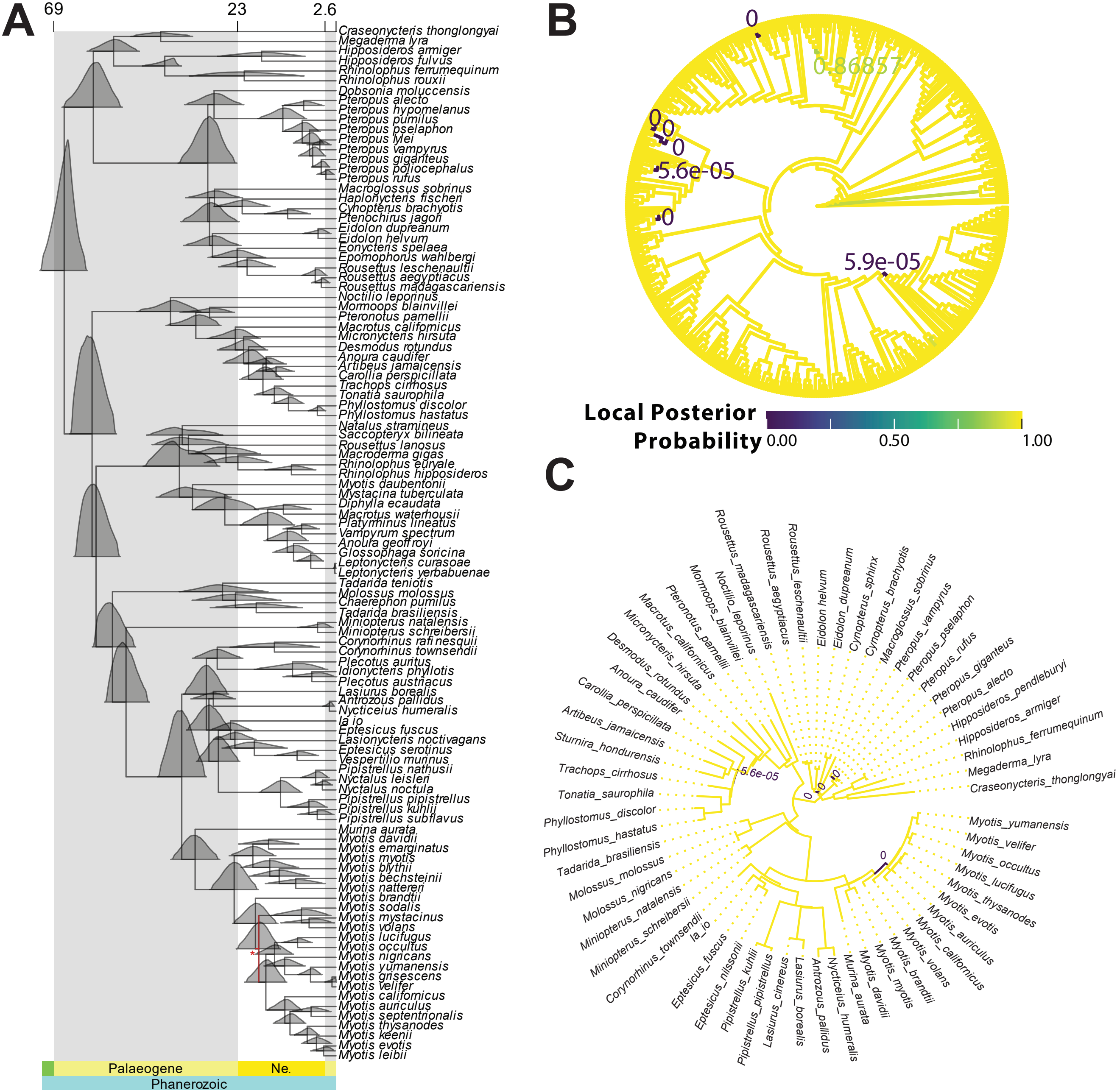

### ExtendedDataFig_3.pdf

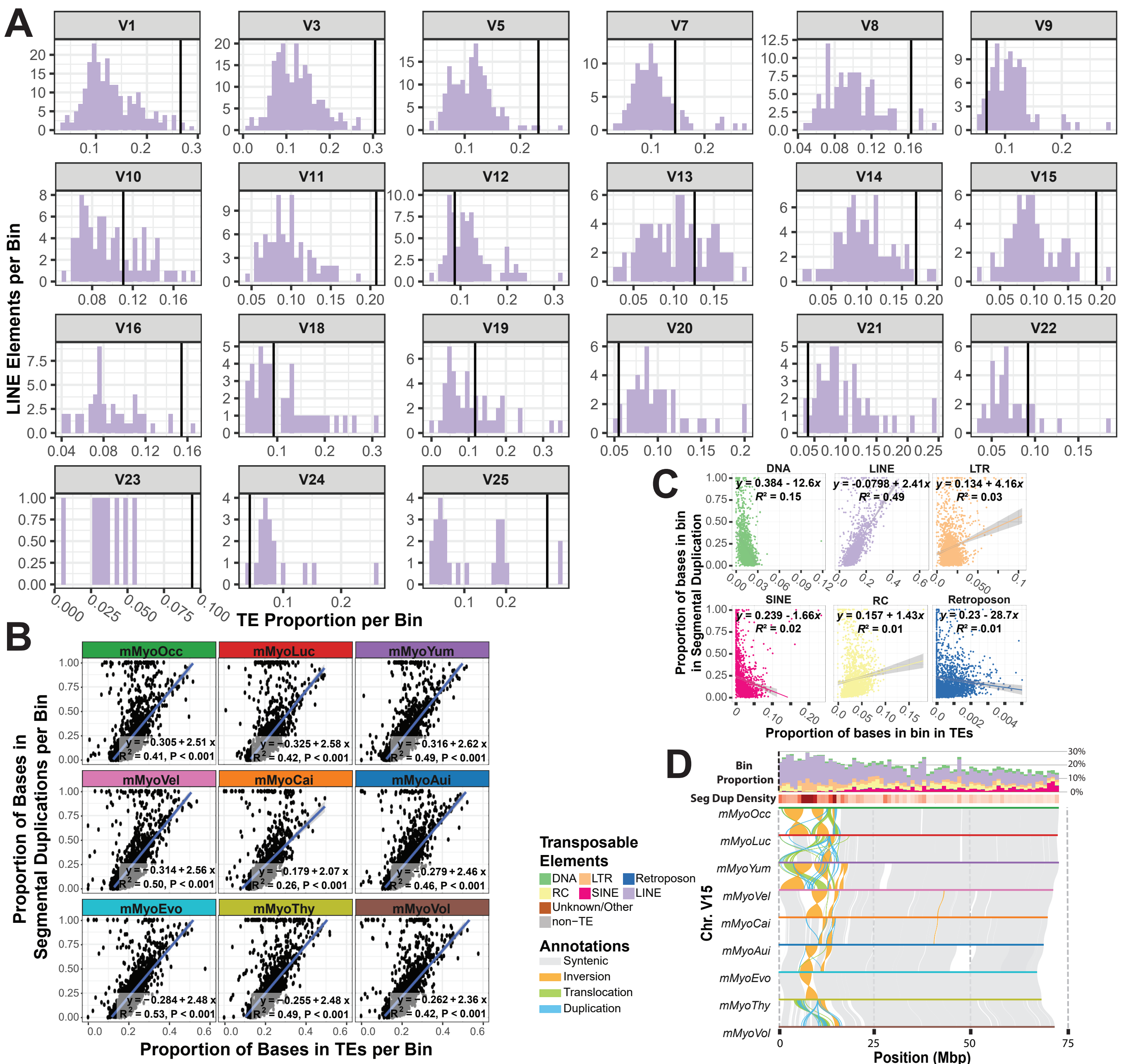

### ExtendedDataFig_4.pdf

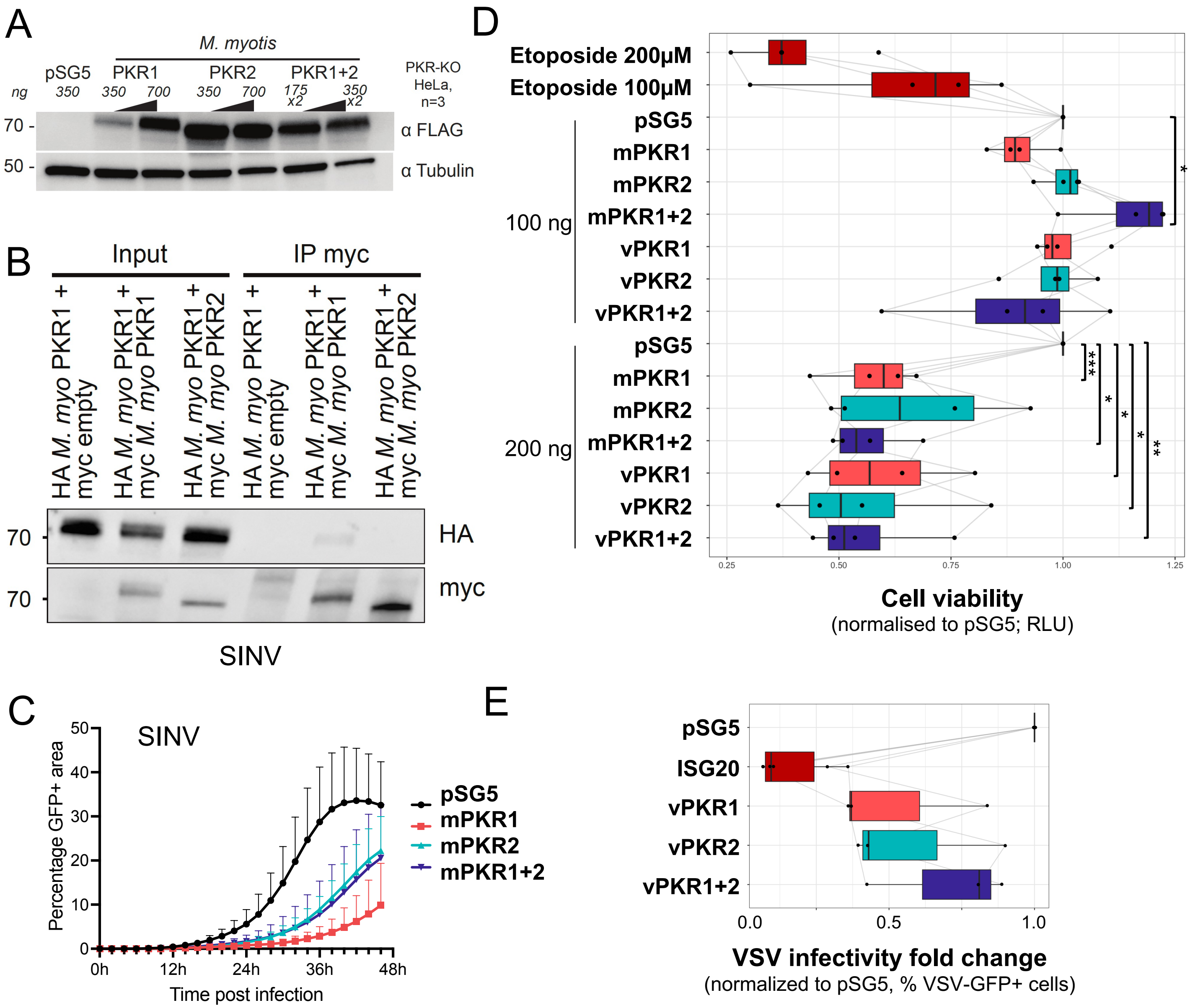

### ExtendedDataFig_5.pdf

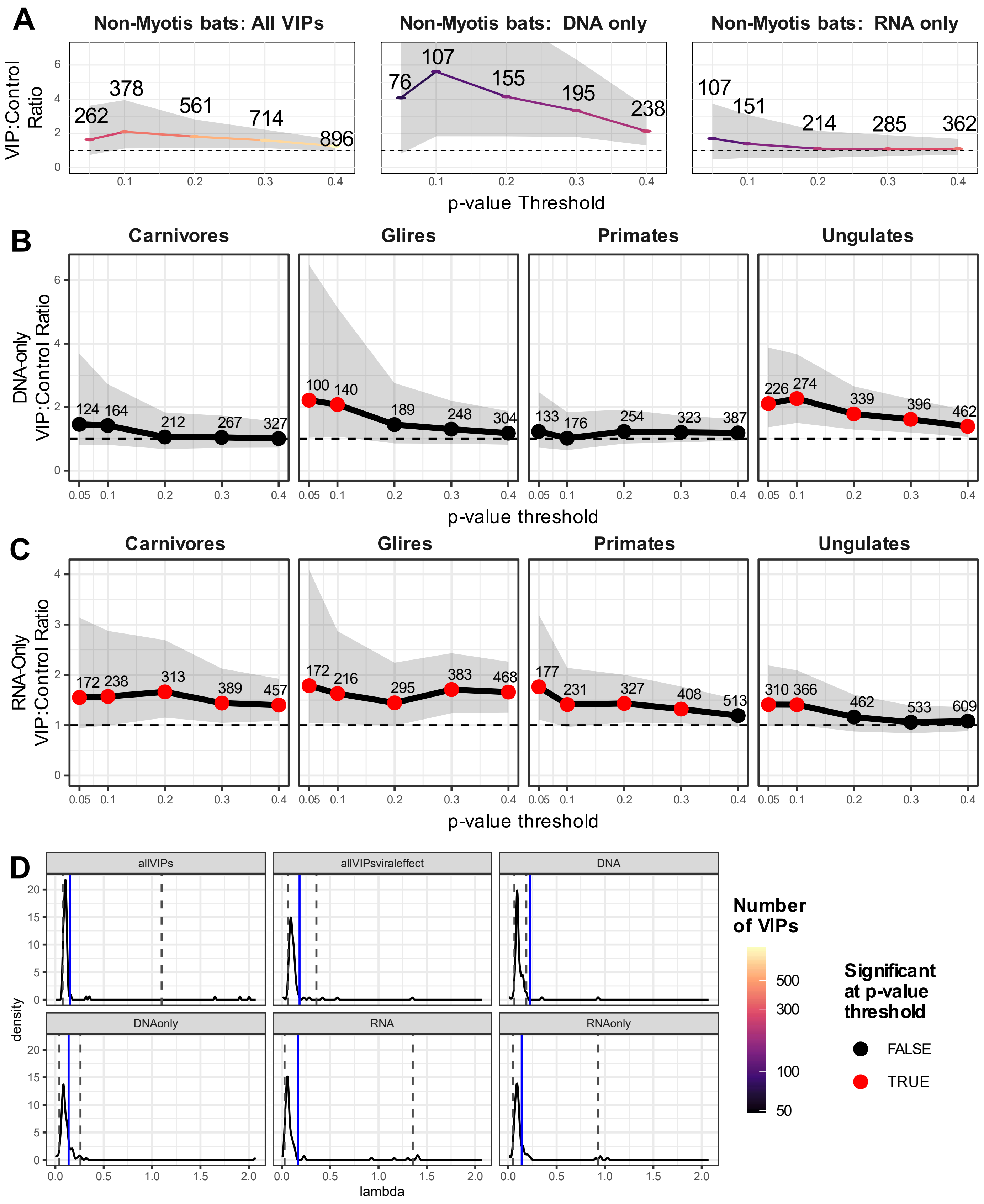

### ExtendedDataFig_6.pdf

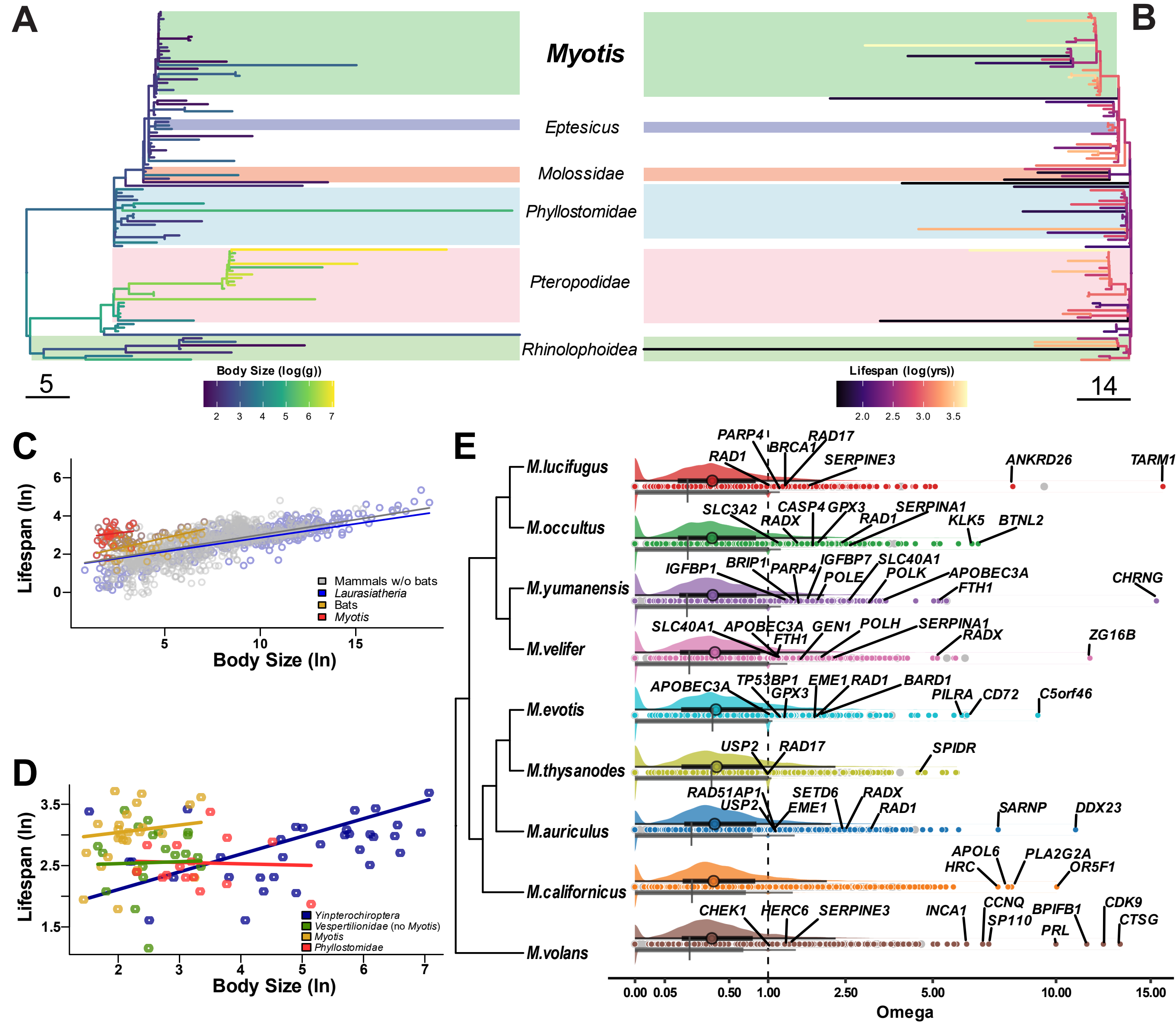

### ExtendedDataFig_7.pdf

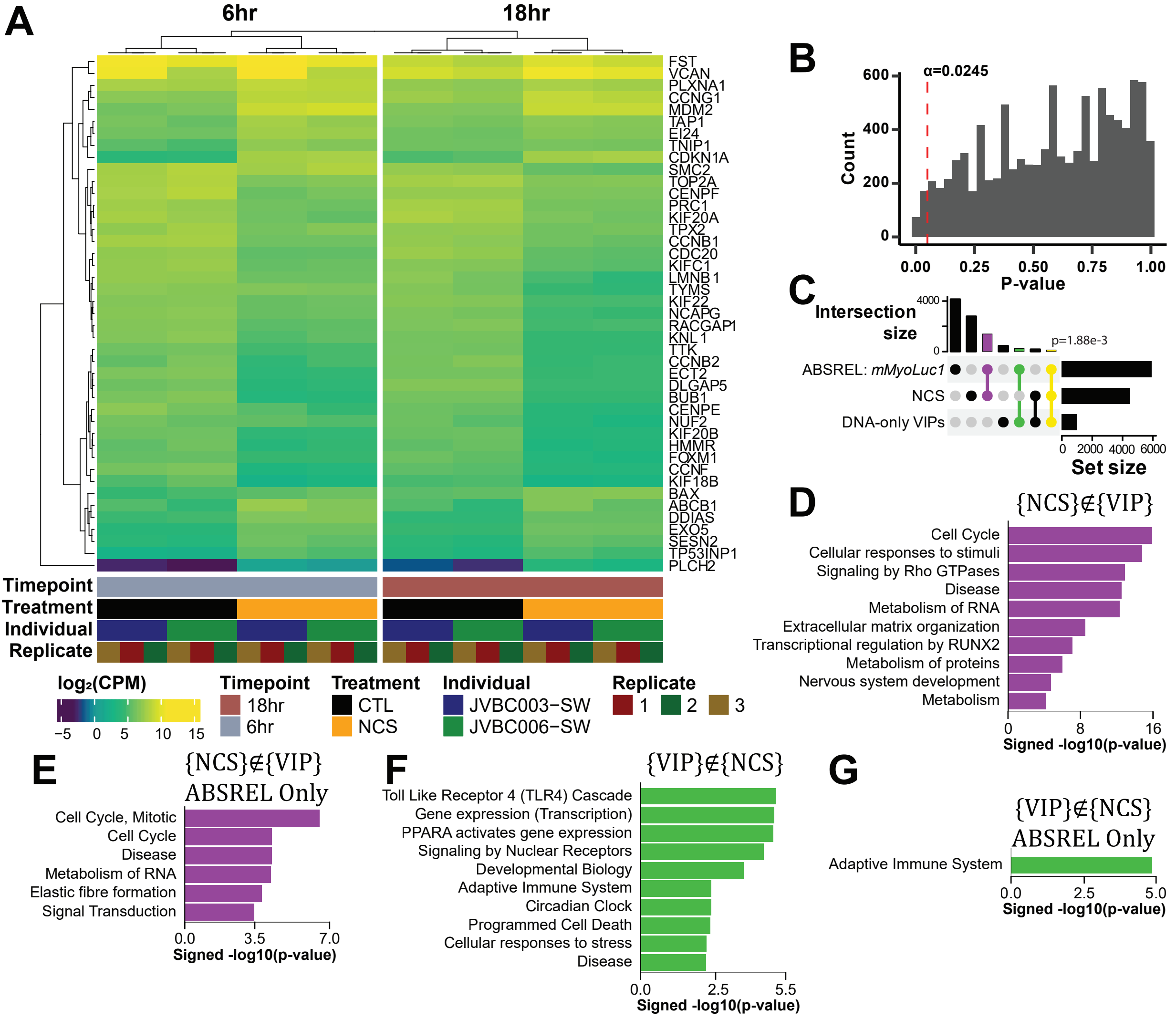
