## Supplementary Information Guide for "Insights into longevity and virus-driven adaptation from *Myotis* bat genomes"

### Affiliations:

### **Table of Contents:**

I. Supplementary Methods
   A. Sample collection and primary cell line generation
   B. RNA extraction, library preparation, and sequencing
   C. DNA extraction, library preparation, and sequencing
   D. Contig Assembly, Scaffolding, and Manual Curation
   E. Identification and annotation of repetitive elements
   F. Gene predictions
   G. Gene prediction curation
   H. Mapping PKR exons
   I. Orthologous Gene Alignments
   J. BUSTED-MH
   K. Viral-Interacting Protein (VIP) Selection Enrichment
   L. Neocarzinostatin Treatment and RNA-seq
II. Supplementary Notes
    A. On Peto’s Paradox and cancer prevalence estimation in bats
    B. DNA & RNA antiviral immunity
    C. Pleiotropy between VIPs and Hallmarks of Aging
III. Supplementary References

### I. **Supplementary Methods**

##### Sample collection and primary cell line generation

For *M. volans, M. occultus, M. auriculus,* and *M. californicus*, two 3-mm wing punch biopsies were taken from the left and right plagiopatagium of each donor individual and placed in a live cell collection media consisting of DMEM/F12 (Gibco) supplemented with 15mM HEPES (Gibco), 20% FBS (Gibco), and 0.2% Primocin (Invivogen)^96–98^. Wing punches were then brought back to a cell culture facility in Berkeley, where they were used to generate cell lines as previously described^96–98^. Additional cell lines for *M. lucifugus, M. velifer, M. yumanensis, M. evotis,* and *M. thysanodes* were similarly collected and generated.

Cell lines for the *M. evotis* and *M. thysanodes* genomes were generously provided by Richard Miller. Cell lines for functional work in *Rousettus langosus*, *Pteropus rodrigensis*, and *Eidolon helvum* were provided by the San Diego Frozen Zoo.

##### RNA extraction, library preparation, and sequencing

To assist our annotation efforts, we generated mRNA-seq for 7 of the species sequenced *de novo* in this study. For *M. velifer*, samples of heart, brain, kidneys, lungs, pancreas, and testis collected from the donor individual were provided to Dovetail Genomics (CA, USA) for mRNA-seq library preparation and sequencing. For *M. occultus, M. thysanodes, M. evotis, M. volans, M. auriculus,* and *M. californicus*, we extracted total RNA from the same cell lines used for the genomic libraries using the RNEasy Plus Mini Kit (Qiagen). Next, we generated rRNA-depleted total RNA-seq libraries using the NEBNext rRNA Depletion Kit v2 and Ultra II Directional RNA Library Prep Kits. RNA and libraries were quality controlled on an Agillent Bioanalyzer using the RNA 6000 Nano and DNA High Sensitivity assays, respectively. Samples were sequenced on to 100M 150PE reads per sample using the Novoseq platform (Novogene). For *M. lucifugus*, we used the following published RNA-seq data on NCBI SRA generated using poly-A selection and paired-end sequencing: SRR6793287, SRR6793288, SRR6793289, SRR6793290, SRR6793291, SRR6793292, SRR6793293, SRR6793294, SRR6793295, SRR6793296, SRR6793297, SRR6793298, SRR6793299, SRR6793300, SRR6793301, SRR7064951, SRR10512805, SRR10512806, SRR10512807, SRR10512808, SRR10512809, SRR10512818, SRR10512829, SRR10512840, SRR10512845, SRR10512846, SRR10512847, SRR10512848, SRR10512849, SRR10512850, SRR10512851, SRR10512852, SRR10083333, SRR10083334, SRR10083335, SRR10083336, SRR10083337, SRR10083338, SRR10083339, SRR10083340, SRR10083351, SRR10083352, SRR1916825, SRR1916826, SRR1916827, SRR1916830, SRR1916832, SRR1916834, SRR1916836, SRR1916839, SRR1916841, SRR1916842, SRR18761564, SRR18761566, SRR18761568, SRR18761571, SRR18761573, SRR18761563, SRR18761565, SRR18761567, SRR18761569, SRR18761570, SRR18761572, SRR18761574, SRR1270869, SRR1270914, SRR1270919, SRR1270921, SRR1270922, SRR1270923, SRR4249979, SRR4249988, SRR5676382, SRR5676383, SRR5676395, SRR5676396, SRR5676402, SRR1869462, and SRR1013468.

##### DNA extraction, library preparation, and sequencing

For 6 genomes (*M. evotis, M. thysanodes, M. volans, M. occultus, M. auriculus,* and *M. californicus*) DNA was extracted from primary cell lines expanded from 3M cells at Passage 2-4 to approximately 40M cells per line using a Circulomics BigDNA CCB kit following the UHMW protocol for cells. DNA from *M. lucifugus* was extracted from flash-frozen tissue by the Genetics Lab of the California Department of Fish and Wildlife. PacBio HiFi libraries were generated and sequenced on a Sequel II (PacBio) by the Functional Genomics Core at the University of California, Berkeley. For cell-line-derived genomes, Hi-C libraries for these genomes were generated from 1M cells at Passage 3 using the OmniC for Illumina kit (Dovetail genomics); libraries were submitted for quality control and sequencing on the Illumina NovaSeq platform (Novogene). For the *M. velifer* genomes, DNA was extracted from flash-frozen tissues, and all DNA extraction, library prep, and sequencing was completed by Dovetail Genomics following standard protocols. For *M. lucifugus*, a previously published Hi-C dataset from 4 pooled individuals was used for scaffolding^99,100^.

The PacBio reads were processed using SMRTTools (v6.0.0-1, PacBio) to generate the circular consensus sequences using the settings “*--minPasses=3 --minRQ=0.99*”. Hi-C reads were processed using trimmomatic^101^ (v0.35-6) to remove adapter sequences and low-quality bases using the settings “*ILLUMINACLIP:data/trimmomatic-adapters/TruSeq3-PE-2.fa:2:40:15 SLIDINGWINDOW:5:20”*.

##### Contig Assembly, Scaffolding, and Manual Curation

To generate the primary contig assemblies, we used hifiasm^102,103^ (v0.14-hd174df1_0) in Hi-C mode, providing both the CCS reads and the trimmed Hi-C reads as input, and purging duplicates using the -l2 option. For our reference genomes, we proceeded with the primary contig assembly (“{species}.asm.hic.p_ctg.gfa”).

All reference genomes were scaffolded with YAHS^104^ (v1.1a.1s) and the Hi-C datasets. Dovetail Omni-C data were processed and mapped to the genome following the manufacturer's instructions using bwa^105,106^ (v0.7.17-h5bf99c6_8), pairtools^107^ (v0.3.0-py37hb9c2fc3_5), and samtools^108^ (v1.12-h9aed4be_1). YAHS was run using both default settings as well as with --no-contig-ec; after comparing the outputs, we proceeded with the --no-contig-ec version for our final assemblies.

To finalize the assemblies, we performed manual curation using PreTextView^109^ and the Rapid Curation toolkit^110^ (version ff964069). The X chromosomes were identified based on size, synteny across genomes, and half-coverage observed in XY genomes; putative Y chromosomes were similarly identified in XY genomes. Mitochondrial genomes were identified and removed from the final assembly by running *mitohifi**^111^* (v3.0) in contig mode on the assembly and removing all scaffolds identified as mitogenomes. The consensus mitogenome from *mitohifi* was designated as the representative mitogenome for the assembly after manual curation. Finally, to eliminate spurious duplicates, we used FunAnnotate^112^ (v1.8.15) and the “clean” function to identify and remove any remaining scaffolds with 90% identical to a larger scaffold.

##### Identification and annotation of repetitive elements

We used RepeatMasker^113^ (version 4.0.7-open) to annotate repetitive elements in our genomes. We first ran RepeatMasker using a curated database of transposable elements from 249 mammalian species^82,114^ (David Ray, pers. comm.) and the settings “*-engine ncbi -s -noisy -xsmall*” followed by a second run using RepeatModeler^115^ and RepeatMasker to identify *de novo* repeats missing from the curated database. All repeats were then soft-masked in all genomes. To assess the repeat landscape, we calculated the summary of divergence from the repeat alignments and created the repeat landscape using auxiliary RepeatMasker scripts (calcDivergenceFromAlign.pl & createRepeatLandscape.pl).

##### Gene predictions

To create optimal gene annotations, we combined *ab initio* gene predictions; orthology inferences; and transcriptomic evidence for a total-evidence dataset facilitated using FunAnnotate^112,116^ with manual interventions. To generate high-quality orthology-based evidence, we downloaded the UNIPARC database^117^ of genes present in all Chiropteran genomes and mapped these proteins to our genomes using miniprot^118^ (v 0.6-r194-dirty). We assembled our transcriptome data into transcripts using TRINITY^119^ (v 2.13.2), and mapped these transcripts to our genomes using minimap2^73^ (v 2.24).

Next, we ran BUSCO^17,120^ (version 5.4.3) using the “eutheria_odb10” database and AUGUSTUS^121^ to identify BUSCO orthologs in our genomes. GFFs describing the gene structure of single-copy BUSCO orthologs was then used by FunAnnotate to train SNAP^122^ and GlimmerHMM^123^ (v 3.0.4) prior to gene prediction. GeneMark-ES^124^ (v 4.72) was run using its self-trained model. AUGUSTUS^125^ (v 3.4) was run using a previously-generated model jointly trained on 6 high-quality bat genome assemblies^82^ and supplemented with protein and transcriptome hints generated by FunAnnotate from the UNIPARC and Trinity datasets.

To leverage high-quality annotations from other genomes, we used TOGA^126^ (version 1.0.1) to generate gene annotations for each of our species, using inference from hg38 annotations. TOGA outputs a table of genes (“reg” genes) associated with the projected transcripts from the reference genomes, and a BED file describing the CDS structure of these projected transcripts. To generate a final GFF file summarizing these data, we converted the original BED file to a GFF file; removed the erroneous “Gene” level attributes; and added in new “Gene” entries describing the TOGA-designated genes, modifying the “Parent” attributes of the mRNAs to refer to the correct parent gene. Transcript projections that were not associated with a TOGA gene designation were then dropped.

Finally, we used LiftOff^127^ (v1.6.3) to lift over annotations from the *Myotis myotis* genome (mMyoMyo1.0_primary^82^). Using BUSCO and manual curation, we assessed both the original GenBank (GCF_014108235.1) and NCBI RefSeq (GCA_014108235.1) annotations, and selected the NCBI RefSeq annotation, as it had slightly improved BUSCO scoring and less erroneous intron-exon junctions at select genes. We removed all non-protein-coding genes from the initial GFF file, then ran LiftOff using the settings “ -exclude_partial -polish -cds”.

We evaluated each line of evidence by assessing their completeness using BUSCO and comparing the completeness score to the total number of predicted genes. We found that SNAP and GLIMMERHMM performed the poorest for gene annotations, with both the lowest BUSCO scores and the highest number of low-quality predictions. The miniprot-UniParc and TOGA-hg38 datasets generated the highest quality gene prediction datasets, with near-complete BUSCO scores and reduced low-quality protein predictions.

##### Gene prediction curation

We used EvidenceModeler^128^ (version 2.0) to generate an initial consensus gene set using only the best lines of evidence (AUGUSTUS, weight 2; high quality AUGUSTUS, weight 5; TOGA-hg38, weight 12; miniprot-UniParc, weight 5; and LiftOff-mMyoMyo1, weight 5) with hints from protein orthology (miniprot-UniParc, weight 6) and RNA-seq (TRINITY, weight 5) for alternative splicing. By default, EvidenceModeler does not consider genes that are located within intronic regions of other genes. To restore these genes, we intersected the EvidenceModeler consensus gene GFF with the TOGA-hg38 GFF to identify which genes were present in intronic regions and omitted from EvidenceModeler; these genes were then added back to the EvidenceModeler gene set.

To eliminate remaining spurious predictions, we cross-referenced our gene annotations against the SwissProt^129^ database using DIAMOND^130^ (v. 2.1.4) with settings “--*ultra-sensitive --outfmt 6 qseqid bitscore sseqid pident length mismatch gapopen qlen qstart qend slen sstart send ppos evalue --max-target-seqs 1 --evalue 1e-10* ”. We kept all genes that matched a protein on SwissProt with at least 80% identity, matched over 50% of the target sequence, and coded for at least 50 amino acids. Of the remaining genes, we kept them only if they contained both a start and stop codon with no internal stop codons.

Finally, we further curated our annotations by putting the EVM and TOGA gene predictions in competition with each other when they both annotated the same locus, but with different overlapping or neighboring annotations. In such cases, one of the gene annotations is likely closer to the truth. To determine which, we compared EVM and TOGA gene models with their closest human gene BLAST hits. Only proteins with a BLAST match to a human Ensembl v99 annotation with the lowest E-values below 0.001 were considered. These human homologs were used as a reference for curation as they are well-defined and characterized. We observed that occasionally, either the EVM or TOGA model predicted a transcript much longer than their human closest homolog. Closer inspection revealed that such cases represent artifactual mergers of neighboring genes during the annotation process, clearly visible from the fact that they map to two distinct human homologs in succession. Such cases were resolved by choosing the annotations (between EVM and TOGA) that were not affected by the artificial merger. We further observed a specific class of mergers between neighboring, segmentally duplicated genes, with the resulting annotations representing chimeric mixes of exons from the duplicates. In such cases we selected the annotations that clearly stayed within the boundaries of the separate duplicates, as identified by mapping to the closest human homolog. For the remaining annotations where both TOGA and EVM both mapped to a single human homolog throughout their entire length, we selected the most complete annotation that was closest in length to the human homolog.

##### Mapping PKR exons

We further validated the annotations for the PKR locus by re-aligning the primary *M. velifer* coding sequence back to the nine nearctic *Myotis* reference genomes, as well as a non-*Myotis* outgroup, *Pipistrellus pygmaeus*, and the genome haplotypes for each of these species. Because the presence of two copies makes this task challenging for most aligners, we independently aligned the *M. velifer* reference PKR sequence to sequential sections of each genome in 50kb search regions surrounding the known loci in each genome. This alignment search was conducted for 5 regions upstream (250 kb) and 5 regions downstream (250 kb) of the known loci. In species with two known copies, the location of each copy was included in a separate search region. This was to prevent erroneous merging or loss of exons. These regions were retrieved using bedtools getfasta^131^ and alignment was performed using miniprot^118^. Miniprot settings were optimized to retain secondary alignments (-p 0 -n 1 –outsc=0.0 –outc=0.0) and find known exons with accurate boundaries (-J 18 -F 21 -O 15 -L 10). The resulting gff file was converted to bed format using AGAT^132^, sequences retrieved with bedtools getfasta, and a custom script used to remove identical duplicates. Finally, all sequences were aligned with MACSE v2.07^133^. We used BISER^19,134^ to confirm the presence of segmental duplications in these regions.

##### Western blot

We assessed for the steady state protein expression of *M. myotis* Flag-PKRs after transfection of 350 ng or 700 ng total of DNA plasmids encoding: PKR1 alone, PKR2 alone or both PKR1 and PKR2 (175 ng of each and 350 ng of each, respectively) in PKR-KO Hela cells^135^. Briefly, cells were re-suspended and lysed in ice-cold RIPA buffer (50 mM Tris pH8, 150 mM NaCl, 2 mM EDTA, 0.5% NP40) with protease inhibitor cocktail (Roche) and sonicated. 20 µL of the clarified fraction was denatured with 5 µL of 6x Laemmli buffer at 95°C for 5 min and loaded into 4-20% BioRad Criterion TGX Stain-Free precast gel. The wet transfer into a PVDF membrane was executed overnight at 4°C. The membranes were blocked in a 1xTBS-T buffer (Tris HCl 50 mM pH8, NaCl 30 mM, 0.05% of Tween 20) containing 10% powder milk, and were incubated for 1h. The membranes were incubated with primary mouse anti-Flag (Sigma F3165) and anti-Tubulin (Sigma T5168) antibodies and secondary anti-Mouse IgG-Peroxidase conjugated (Sigma A9044). Detection was made using the Chemidoc Imaging System (BioRad) with SuperSignal West Pico Chemiluminescent Substrate (ThermoFisher Scientific).

##### Luciferase reporter assays

Luciferase reporter assays were carried out to investigate whether the two PKR paralogs have synergistic, additive or dominant negative effect in translation shutdown. 25,000 PKR-KO cells were seeded in 24-well plates. The next day, cells were transfected with 350 ng of empty pSG5, 350 ng of PKR1, 350 ng of PKR2, or 175 ng of PKR1+ 175 ng of PKR2, with additional 50 ng of FFLuc firefly luciferase reporter plasmid per well. Sterile alpha motif domain-containing proteins 9L (SAMD9L gain-of-function mutant) was used as a positive control of translational repression^72^. 24 hours post transfection, cells were washed with PBS, lysed by a 5× reporter lysis buffer (Promega) and incubated overnight at -20°C. Cells were then collected and 100 μl of the luciferase substrate (Promega) was added to 20 μl of the lysis supernatant. Alternatively, cells were lysed using BrightGlow Lysis Reagent (Promega E2620). The relative luminescence units (RLUs) proportional to the translation of the luciferase reporter were immediately quantified with LUMIstar Omega microplate reader optima (BMG Labtech). All luciferase assays were conducted in technical duplicates in at least five independent experiments. Fold change results were normalized to the empty pSG5 condition within each independent experiment.

##### Orthologous Gene Alignments

Phylogeny and selection analyses described in this manuscript rely on high-quality alignments of bat orthologous coding sequences. To first find and align orthologous *Myotis* genes to the greatest extent possible, we first complemented the gene annotations described above with likely missing annotations that could still be found through BLAT homology searches. Missing gene annotations are always expected in non-model species genomes and reflect a feature of annotation pipelines in general, not an artifactual issue. For example if the first coding exon of a gene falls into a small local assembly gap, the lack of a start codon may prevent the trigger of a CDS annotation, or may lead to the clearly incomplete CDS being subsequently filtered out. Similarly, erroneous indels representing sequencing errors may interrupt coding reading frames. Genes with missing annotations can still be detected in assemblies through classic BLAST or BLAT homology searches, and then aligned with their annotated orthologs from other species. To align orthologous *Myotis* genes from ten species (those sequenced here plus *Myotis myotis* and *M. yumanensis*), we first decided to use *Myotis velifer* as the *Myotis* species of reference, since the RNA-seq data we used was generated with *M. velifer* tissues.

We first looked for missing homologs of *M. velifer* genes in the other *Myotis* genomes by blatting *M. velifer* CDS to the other *Myotis* assemblies (BLAT command line including non-default options -q=dnax -t=dnax -fine) to find matches outside of already annotated genomic segments. When multiple velifer CDS matched to the same locus with multiple overlapping homologous BLAT matches, we selected the match with the highest number of identical nucleotides. The remaining matching BLAT sequences were further considered if they spanned at least 50% of the velifer CDS, and included 100 codons or more. BLAT matches including stop codons were removed. This process added 1,837 putative CDS to consider for orthologous alignments for *M. auriculus*, 1,785 for *M. californicus*, 1,796 for *M. evotis*, 1,505 for *M. lucifugus*, 3,234 for *M. myotis*, 1,826 for *M. occultus*, 1,822 for *M. thysanodes*, 1,800 for *M. volans* and 1,729 for *M. yumanensis*. The correct reading frames for these putative CDS were then determined by aligning to the velifer CDS that generated the initial match with MACSE v2. MACSE has the crucial advantage over other aligners of being able to repair broken reading frames due to sequencing indel errors or erroneous gene annotations. At this stage, we restricted any further analysis to those velifer CDS with human homologs (BLASTP E-value<0.001 with at least one human canonical protein-coding gene from Ensembl). One-to-one orthologs with the 23,030 remaining velifer CDS in other *Myotis* species were then determined using Orthofinder v2.5.4^136^. The sequences of each group of ortholog were then aligned with MACSE v2^133^ with default settings. The resulting CDS with potentially repaired reading frames were then checked with PREQUAL^137^ to exclude sequencing errors and erroneous inclusion of non-homologous segments in annotations. The remaining parts of orthologous sequences that passed PREQUAL filtering were then aligned again using MACSE v2 with default settings. The first round of alignment with MACSE ensures that we do not remove portions of CDS that look like they have no homology and would thus be removed by PREQUAL, just because of frameshifts that are easy to repair first with MACSE. The second round of MACSE is to align the remaining codons once PREQUAL has removed erroneous portions of CDS that could have otherwise disturbed the alignment process. We further masked (i.e. replaced with indels) codons near indels with putative alignment errors as described in Bowman et al.^138^. Of the 23,030 initial *M. velifer* CDSs, this process resulted in 21,756 alignments with at least one ortholog in another *Myotis* species.

We also aligned pan-Chiroptera orthologs from 47 non-*Myotis* genomes publicly available on NCBI at the time of analysis, to test the generality of our observations to all bats. We used the same strategy described above to complement *Myotis* gene annotations with BLAT matches, but this time blatting velifer CDS on non-Myotis assemblies (with -q=dnax -t=dnax -fine again) to find all the potential orthologs in the non-Myotis assemblies. We previously found that because BLAT represents a first filter to include only portions of homologous CDS with good local similarity, using BLAT matches results in higher quality alignments of orthologs than using existing gene annotations of disparate qualities that too often include non-homologous portions of introns among other issues^138,139^. As before with only *Myotis* species, we recovered putative one-to-one orthologs with Orthofinder. This process resulted in the alignment (as previously described with two rounds of MACSE and PREQUAL in the middle) of 19,009 orthologous CDS with at least one non-*Myotis* orthologous CDS.

To test whether the patterns of virus-driven adaptation observed in bats are unique across mammals, we also prepared four more datasets of 70 primate orthologous CDS alignments, 138 euungulate alignments, 127 glire alignments, and 82 carnivora alignments (see **Table S2** for the species and their respective assemblies used), the orders with the most species represented in our dataset. We used the same pipeline as the one used to align 47 pan-Chiroptera species as described above, except that instead of starting from velifer CDS, we started from human Ensembl v109^140^ CDS (the longest isoform available in each case) for primates, *Mus musculus* Ensembl v109 longest CDS for glires, *Canis familiaris* Ensembl v109 longest CDS for carnivores, and *Bos taurus* Ensembl v109 longest CDS for euungulates . These species were chosen for the very high quality of their gene annotations.

##### BUSTED-MH

The original BUSTED test estimates for a given gene the proportion of codons that have evolved under positive selection, with dN/dS>1, summed over all the branches of a given tree, regardless of the branch and regardless of the codons in a multi-species alignment. The version of BUSTED we used, BUSTED-MH, includes two crucial improvements over the original BUSTED that make it much less likely to generate false positive inferences of positive selection, albeit at the cost of becoming a very conservative test. First, BUSTED-MH takes synonymous substitution rate variation into account, which prevents mistaking cases where dN/dS is greater than one just because dS is low, with cases where dN/dS is greater than one because positive selection actually increased dN. Second, BUSTED-MH takes complex substitutions that simultaneously involve more than one nucleotide into account in its likelihood models. This prevents attributing positive selection to cases where dN/dS is greater than one where instead a complex substitution changed multiple amino acids in a single event. BUSTED-MH has been shown to strongly reduce the rate of false positives that typically plague dN/dS-based tests of positive selection^141^.

##### Viral-Interacting Protein (VIP) selection enrichment

To determine if *Myotis* and other bats are enriched for adaptation at Virus Interacting Proteins (VIPs), we conducted a test comparing levels of adaptation, inferred by BUSTED, in sets of VIP genes compared to matched control genes. Sets of control genes were resampled in a bootstrap procedure (https://github.com/DavidPierreEnard/Gene_Set_Enrichment_Pipeline) to generate 95% confidence intervals for sets of genes at progressively smaller BUSTED p-value thresholds^24,26,32,59^. When VIPs are subject to greater levels of positive selection than expected relative to the sets of matched control genes, we expect a pattern in which the high p-value thresholds show weaker enrichment but smaller confidence intervals, because more genes are used in these calculations. As the p-value threshold gets smaller, the signal of enrichment is expected to get stronger but at the expense of larger confidence intervals.

We generated five sets of VIP genes: A set of all VIP genes with aligned orthologs from at least five species in the tested clade (Nearctic *Myotis* or pan-Chiroptera without *Myotis*); a set of VIP genes with known pro- and/or anti-viral activity; a set of VIP genes with no known pro- and/or anti-viral activity; a set of VIP genes that interact only with DNA viruses (and do not interact with RNA viruses); and a set of VIP genes that interact only with RNA viruses (and do not interact with DNA viruses). We divide VIPs into DNA-only and RNA-only because it has been previously found that the DNA/RNA virus distinction in humans is evolutionary relevant - RNA viruses have driven most adaptation in humans^32^, and most zoonoses are caused by RNA viruses. Because both the number of species and genes included, as well as their level of homology, influences the power of these tests we also tested the influence of the stringency of gene choice by generating a separate set of genes for the pan-Chiroptera analyses that included only genes with aligned orthologs in at least two thirds of the non-*Myotis* species. Analyses using this more limited set of genes show the same result in terms of enrichment of adaptation in VIP genes and comparing DNA VIPs and RNA VIPs, showing that the observed patterns are valid across bats regardless of the stringency of homology.

The bootstrap procedure matches a tested gene set of interest such as VIPs with sets of control genes (non-VIPs when testing VIPs) that have the same average values as the set of interest for multiple potential confounding factors that could explain differences in adaptation instead of interactions with viruses. For example, if the level of gene mRNA expression has an influence on the rate of adaptation, we then need to match VIPs with control sets of non-VIPs that collectively have the same average expression as VIPs. For each group of tested VIPs we build 1,000 control sets with randomly sampled non-VIPs according to a matching procedure described in Enard & Petrov 2020^59,71^. We match the following factors between VIPs and non-VIPs, for all tested groups of species:

25. The length of the aligned CDS.
26. The overall CDS GC content in each orthologous alignment.
27. The GC content at aligned codons’ position 1, 2 and 3 separately.
28. The number of species with a one to one ortholog out of all the species included in an alignment, where
    species with no ortholog are represented by gaps the whole length of the alignment.
29. The number of species with an ortholog at least 90% of the length of the species of reference (Myotis velifer in bats, human in primates, etc; see above).
30. The overall proportion of each orthologous alignment made of indels.
31. The three synonymous rates of evolution estimated by the likelihood model of HyPhy.
32. The proportions of codons that fall in the three latter synonymous rates.
33. Average human mRNA expression in 53 GTEx v7 tissues^142^, in log_2_ of Transcripts Per Million (TPM).
34. Lymphocyte human mRNA expression from GTEx v7, in log_2_ of TPM.
35. Testis human mRNA expression from GTEx v7, in log_2_ of TPM.
36. mRNA expression in log_2_ of TPM for six separate *Myotis velifer* tissues: heart, brain, kidneys, lungs, pancreas, and testis.
37. The number in log_2_ of protein-protein interactions (PPIs) in the human protein interaction network^143^.
38. The proportion of genes that are immune genes according to Gene Ontology annotations of the closest human homolog including Gene Ontology terms GO:0002376 (immune system process), GO:0006952 (defense response), and/or GO:0006955 (immune response) as of summer 2021^144^.
39. The proportion of housekeeping genes defined as genes with stable expression across many human tissues, listed in Eisenberg & Levanon^145^.
40. The overall dN/dS ratio estimated by BUSTED for the orthologous CDS alignments.

We match the overall dN/dS between VIPs and control non-VIPs to account for an important issue of dN/dS tests: dN/dS-based tests tend to lose statistical power to detect positive selection in CDS alignments with higher selective constraint^146^. The amount of positively selected sites being equal, positive selection tests based on dN/dS tend to have lower statistical power and tend to generate more false negative results when the rest of the coding sequence is more highly constrained. VIPs tend to be much more strongly constrained than non-VIPs^24^, which gives a strong, unfair statistical disadvantage to VIPs when testing positive selection with BUSTED or other HyPhy tests. We limit this issue by matching VIPs and control non-VIPs for dN/dS. Thus, VIPs have an excess of adaptation compared to non-VIPs when they have a balance of the same total amount of non-synonymous changes more tilted towards advantageous rather than neutral amino acid changes. In this case non-VIPs still have less constraint (more neutral changes) than VIPs, and thus still more power to detect positive selection, but not to an extent as severe and unfair as if we did not match the overall dN/dS^24^. In the case where VIPs do not have an excess of adaptation, then they have the same balance of advantageous and neutral amino acid changes resulting in the same overall dN/dS. This is the case of RNA VIPs in bats in this study; this internal negative control shows that the matching of dN/dS works as intended.

##### Neocarzinostatin treatment and RNA-seq

Primary skin fibroblasts from two individuals of *M. lucifugus* were treated in triplicate with either 100 nM neocarzinostatin (Sigma-Aldrich) or an MES vehicle control (Sigma-Aldrich). Samples were continuously treated for either 6 or 18 hours in two distinct batches. After treatment, wells were gently rinsed twice with ice-cold DPBS prior to *in situ* lysis using Buffer RLT plus (Qiagen) supplemented with 10 uM DTT; RNA was then extracted using the RNeasy Plus Mini Kit (Qiagen) following manufacturer’s instructions. Samples were submitted for library prep and Illumina 150-PE sequencing using Novogene.

All RNA-seq data was pre-processed as described earlier and mapped using STAR^147^ (v. 2.7.6) and our custom *mMyoLuc1* annotations, followed by transcript-level quantification (Transcripts per Million, TPM) using RSEM^148^ (v. 1.3.3). Differential expression analysis was performed using the *edgeR**^149^* package (v4.6.3) in R. As the 6- and 18-hour experiments were run in distinct batches with their own controls, we analyzed each timepoint separately following the quasi-likelihood pipeline using the model “*~Individual+Dose*” and testing the coefficient for Dose.

### II. **Supplementary Notes**

#### On Peto’s Paradox and cancer prevalence estimation in bats

Bats, like other long-lived mammals, are widely remarked as being species resistant to cancer on the basis of their unique life history^3,8,40,150^. However, it is important to note that the limited records for neoplasias in the literature is unlikely to be solely due to any cancer resistance, as records of neoplasias do exist for the limited Yinpterochiropteran species that can be kept in captivity^39,54^. However, it is worthwhile to highlight that while *neoplasias* have been observed in bats, *malignant* tumors have not yet been observed^39,54^, in support of the theory of Chiropteran cancer resistance. Unfortunately, the very trait that makes bats appealing for this line of research - their exceptional longevity - also creates an exceptional challenge for longitudinal study; this is further compounded by the fact that the longest-lived bats in the wild, *Myotis* and other Yangochriopterans, fail to thrive in captivity^151^. These facts underscore the necessity of proxy measurements for cancer resistance in these and other species which cannot be studied in captivity, such as measuring somatic mutation rates across lifespan^152^ and the use of *in vitro* models of carcinogenesis to experimentally test the cancer resistance of long-lived bats^153^. The study of patterns and mechanisms of cancer risk and resistance across mammals will continue as survey-scale technologies for detecting cancers without necropsy continue to develop^154^.

#### DNA & RNA Antiviral Immunity

While bats are known to mount diversified immune responses to RNA viruses, our results suggest that the co-evolutionary history between bats, DNA viruses, and RNA viruses has led to different approaches to dealing with these two classes of pathogens. Changes in gene copy number in bats’ innate and adaptive immune systems, such expansions in *MHC-I* and *APOBEC3**^82^*, or the loss of the *PYHIN* gene family^155^, may be an effective defense or adaptation, but insufficient or ineffective against DNA viruses. Alternatively, it is possible that differences in the rates of evolution between DNA and RNA viruses have required similarly distinct similar timescales of genetic evolution in bats: VIPs constrained by host biology, such as DNA repair and replication, have had time to evolve against the slower mutation rate of DNA viruses, while the rapid mutation rate of RNA viruses has required bat genomes to develop rapid gains and losses of new and specialized antiviral genes. Due to the intrinsic difficulties of studying ancient genetic samples, testing these hypotheses with the long-term history of ancient viruses will require significant increases in effort and technological advances^156^, but would provide significant insights into how host-pathogen interactions shape the evolution of life.

#### Pleiotropy between VIPs and Hallmarks of Aging

Consistent with the agonistic pleiotropy hypothesis, many of the genes and pathways highlighted in this study have been found to play vital roles across physiological traits in bats and other species. As a core part of the integrated stress response PKR is not only associated with the innate immune response, but also has been identified as a longevity-associated gene in multiple studies^157–159^. Additionally, two genes under selection in nearctic *Myotis* - *FTH1* and *IGFN1* - have been implicated in functional studies as key hibernation genes^160–162^, viral interacting proteins^163–166^, and as pro-longevity genes^167–169^. Similarly, many DNA VIPs such as *BRCA1/2* and *POLG* are core DNA maintenance genes essential for cancer resistance and longevity^47,170–176^. Such essential genes are generally under increased constraint; however, it is possible that internal pressures, such as DNA transposon activity, and external pressures such as viral infections and other environmental stressors, are shaping the fitness landscape of these essential genes such that novel optimizations become accessible to evolutionary processes^68^. Changes in structural genomic variation also play a crucial role here, such as our findings related to PKR and RNA VIPs in *Myotis*. Many of the overarching pathways we observed under selection, such as inflammation, senescence, and ferroptosis, lie directly at the intersection of aging-related immune processes^82,155,162,167,177–184^. More broadly, our findings on pervasive selection on DNA-only VIPs and the extreme evolution of longevity and cancer risk in other bats support a more general hypothesis that traits such as cancer risk, cellular homeostasis, and antiviral response have evolved in tandem due to pleiotropic selection at coinciding points in bats’ evolutionary history. Further work leveraging model systems such as *Myotis* for functional evolutionary genomics is necessary to disambiguate the underlying selective pressures driving these functional genetic changes in bats and beyond.

### III. **Supplementary References**

104.	Zhou, C., McCarthy, S. A. & Durbin, R. YaHS: yet another Hi-C scaffolding tool. Bioinformatics 39, (2023).

105.	Li, H. & Durbin, R. Fast and accurate short read alignment with Burrows-Wheeler transform. Bioinformatics 25, 1754–1760 (2009).

106.	Li, H. & Durbin, R. Fast and accurate long-read alignment with Burrows-Wheeler transform. Bioinformatics 26, 589–595 (2010).

107.	Open2C et al. Pairtools: From sequencing data to chromosome contacts. PLoS Comput. Biol. 20, e1012164 (2024).

108.	Danecek, P. et al. Twelve years of SAMtools and BCFtools. Gigascience 10, (2021).

109.	Harry, E. PretextView (Paired REad TEXTure Viewer): A desktop application for viewing pretext contact maps. (2022).

110.	Rapid curation. GitLab https://gitlab.com/wtsi-grit/rapid-curation.

111.	Uliano-Silva, M. et al. MitoHiFi: a python pipeline for mitochondrial genome assembly from PacBio high fidelity reads. BMC Bioinformatics 24, 288 (2023).

112.	Palmer, J. M. & Stajich, J. E. Funannotate: Eukaryotic Genome Annotation Pipeline. (Github, 2023).

113.	Smit, A. F. A., Hubley, R. & Green, P. RepeatMasker Open-4.0. 2013--2015. Preprint at (2015).

114.	Zoonomia Consortium. A comparative genomics multitool for scientific discovery and conservation. Nature 587, 240–245 (2020).

115.	Flynn, J. M. et al. RepeatModeler2 for automated genomic discovery of transposable element families. Proc. Natl. Acad. Sci. U. S. A. 117, 9451–9457 (2020).

116.	Palmer, J. M. & Stajich, J. Funannotate v1.8.1: Eukaryotic Genome Annotation. (Zenodo, 2020). doi:10.5281/ZENODO.4054262.

117.	Leinonen, R. et al. UniProt archive. Bioinformatics 20, 3236–3237 (2004).

118.	Li, H. Protein-to-genome alignment with miniprot. Bioinformatics (2023).

127.	Shumate, A. & Salzberg, S. L. Liftoff: accurate mapping of gene annotations. Bioinformatics (2021).

128.	Haas, B. J., Salzberg, S. L., Zhu, W., Pertea, M. & Allen, J. E. Automated eukaryotic gene structure annotation using EVidenceModeler and the Program to Assemble Spliced Alignments. Genome Biol. (2008).

129.	Quintaje, S. B. & Orchard, S. The annotation of both human and mouse kinomes in UniProtKB/Swiss-Prot: one small step in manual annotation, one giant leap for full comprehension of genomes. Mol. Cell. Proteomics (2008).

130.	Buchfink, B., Reuter, K. & Drost, H.-G. Sensitive protein alignments at tree-of-life scale using DIAMOND. Nat. Methods 18, 366–368 (2021).

131.	Quinlan, A. R. BEDTools: The Swiss-Army Tool for Genome Feature Analysis. Curr. Protoc. Bioinformatics 47, 11.12.1–34 (2014).

132.	Dainat, J. AGAT: Another Gff Analysis Toolkit to Handle Annotations in Any GTF/GFF Format. doi:10.5281/zenodo.3552717.

133.	Ranwez, V., Douzery, E. J. P., Cambon, C., Chantret, N. & Delsuc, F. MACSE v2: Toolkit for the Alignment of Coding Sequences Accounting for Frameshifts and Stop Codons. Mol. Biol. Evol. 35, 2582–2584 (2018).

134.	Išerić, H., Alkan, C., Hach, F. & Numanagić, I. Fast characterization of segmental duplication structure in multiple genome assemblies. Algorithms Mol. Biol. (2022).

141.	Lucaci, A. G., Zehr, J. D., Enard, D., Thornton, J. W. & Kosakovsky Pond, S. L. Evolutionary Shortcuts via Multinucleotide Substitutions and Their Impact on Natural Selection Analyses. Mol. Biol. Evol. 40, (2023).

142.	GTEx Consortium. The GTEx Consortium atlas of genetic regulatory effects across human tissues. Science 369, 1318–1330 (2020).

143.	Luisi, P. et al. Recent positive selection has acted on genes encoding proteins with more interactions within the whole human interactome. Genome Biol. Evol. 7, 1141–1154 (2015).

144.	Gene Ontology Consortium. The Gene Ontology resource: enriching a GOld mine. Nucleic Acids Res. 49, D325–D334 (2021).

145.	Eisenberg, E. & Levanon, E. Y. Human housekeeping genes, revisited. Trends Genet. 29, 569–574 (2013).

146.	Kosakovsky Pond, S. L. et al. A random effects branch-site model for detecting episodic diversifying selection. Mol. Biol. Evol. 28, 3033–3043 (2011).

147.	Dobin, A. et al. STAR: ultrafast universal RNA-seq aligner. Bioinformatics 29, 15–21 (2013).

148.	Li, B. & Dewey, C. N. RSEM: accurate transcript quantification from RNA-Seq data with or without a reference genome. BMC Bioinformatics 12, 323 (2011).

149.	Chen, Y., Chen, L., Lun, A. T. L., Baldoni, P. L. & Smyth, G. K. edgeR v4: powerful differential analysis of sequencing data with expanded functionality and improved support for small counts and larger datasets. Nucleic Acids Res. 53, (2025).

150.	Austad, S. N. Methusaleh’s Zoo: How Nature Provides Us with Clues for Extending Human Health Span. J. Comp. Pathol. 142, S10–S21 (2010).

151.	Lollar, A. The Rehabilitation and Captive Care of Insectivorous Bats. Weatherford International: Houston, TX, USA (2018).

164.	Demogines, A., Abraham, J., Choe, H., Farzan, M. & Sawyer, S. L. Dual host-virus arms races shape an essential housekeeping protein. PLoS Biol. 11, e1001571 (2013).
